# Benzylisoquinoline alkaloid production in yeast via norlaudanosoline improves selectivity and yield

**DOI:** 10.1101/2023.05.19.541502

**Authors:** Lauren Narcross, Michael E. Pyne, Kaspar Kevvai, Ka-Hei Siu, John E. Dueber, Vincent J. J. Martin

**Author notes:** Department of Biology, Western University, London, Ontario, Canada (Michael E. Pyne). Lallemand Inc., Montréal, Québec, Canada (Kaspar Kevvai). Department of Chemical and Biological Engineering, Princeton University, Princeton, New Jersey, USA.

## Abstract

The benzylisoquinoline alkaloid (BIA) family of tetrahydroisoquinolines (THIQs) comprises over 2,500 members, including the pharmaceuticals morphine, codeine, and papaverine as well as the antibiotics sanguinarine and chelerythrine used in animal husbandry. Agricultural cultivation can currently supply the demand for the BIAs that accumulate in plants, but broader access to the entire BIA family would open new avenues of research and commercialization. Microbial synthesis presents an attractive option due to cheap feedstock, genetic tractability, and ease of scale-up. Previously we reported titers of the key branch-point BIA (*S*)-reticuline of 4.6 g/L in yeast, which was achieved through leveraging the Ehrlich pathway 2-oxoacid decarboxylase Aro10 to generate the intermediate 4-hydroxyphenylacetaldehyde (4-HPAA). Here, we establish a superior route to (*S*)-reticuline by switching the pathway intermediate from 4-HPAA to 3,4-dihydroxyphenylacetaldehyde (3,4-dHPAA) using human monoamine oxidase A (MAO). The resulting (*S*)-norlaudanosoline route to (*S*)-reticuline synthesis is more selective, resolving prior issues with off-pathway THIQs synthesized due to concerted enzyme promiscuity. The new pathway is also more efficient, enabling titers of 4.8 g/L (*S*)-reticuline while improving yields over 40%, from 17 mg/g sucrose to 24 mg/g sucrose in fed-batch fermentations. Finally, we extend *de novo* (*S*)-reticuline synthesis to dihydrosanguinarine, achieving 635 mg/L dihydrosanguinarine and sanguinarine in fed-batch fermentation the highest reported titer of these BIAs by a factor of 40.

**Highlights:**

- Monoamine oxidase A (MAO) supports high-titer (*S*)-reticuline synthesis in yeast
- MAO route to (*S*)-reticuline improves specificity compared to Aro10 route
- This work represents a 40% increase in highest reported (*S*)-reticuline yield
- 653 mg/L (dihydro-) sanguinarine was produced by extending the pathway

## Introduction

Benzylisoquinoline alkaloids (BIAs) are a large class of plant secondary metabolites in the tetrahydroisoquinoline (THIQ) family with broad applications across human health and agriculture. While some BIAs accumulate to a sufficient degree in plants to allow for commercial-scale production, most do not. A sustainable, scalable source of BIAs would expand access to this valuable class of natural products. One promising option is the introduction of BIA synthesis to a microbial host.

BIA synthesis from simple carbon sources has been established in *Escherichia coli*^1^, *Saccharomyces cerevisiae*^2, 3^, and *Pichia pastoris*^4^. (*S*)-Reticuline is a common target for *de novo* BIA synthesis, as it is the last shared pathway intermediate in the morphine, sanguinarine, and noscapine pathways^5^. To date, the highest reported (*S*)-reticuline titers are 8 mg/L in *P. pastoris*, 0.16 g/L in *E. coli*^6^, and 4.6 g/L in *S. cerevisiae*^7^. The latter titer is noteworthy because it approaches a target set for commercial production of opioids in microbes: 5 g/L^8^.

The committed step of BIA synthesis in plants is the formation of (*S*)-norcoclaurine by norcoclaurine synthase (NCS) through the condensation of dopamine and 4-hydroxyphenylacetaldehyde (4-HPAA), both derivatives of the aromatic amino acid pathway (Figure 1A)^5^. In yeast, dopamine is a non-native metabolite requiring expression of heterologous enzymes for hydroxylation and decarboxylation of L-tyrosine. 4-HPAA is a native yeast metabolite, derived from the tyrosine precursor 4-hydroxyphenylpyruvate (4-HPP) by the 2-oxoacid decarboxylase Aro10 as part of the Ehrlich pathway of amino acid catabolism. While overexpression of *ARO10* is an effective choice for 4-HPAA overproduction in yeast (Figure 1A)^7, 9^, the enzyme is also capable of synthesizing analogous aldehydes from the 2-oxoacid precursors of the L-amino acids phenylalanine, tryptophan, methionine, leucine, isoleucine, and valine^10, 11^. Compounding the effects of this promiscuity, NCS can condense dopamine with a variety of aldehydes in addition to 4-HPAA leading to the synthesis of additional THIQs, which are in turn substrates for BIA *O*- and *N*-methyltransferases^12, 13^. Although this web of overlapping promiscuities can enable the elucidation and establishment of synthetic routes to new compounds^7^, a more selective mechanism for aldehyde synthesis would further improve targeted BIA production in yeast.

**Figure 1.**
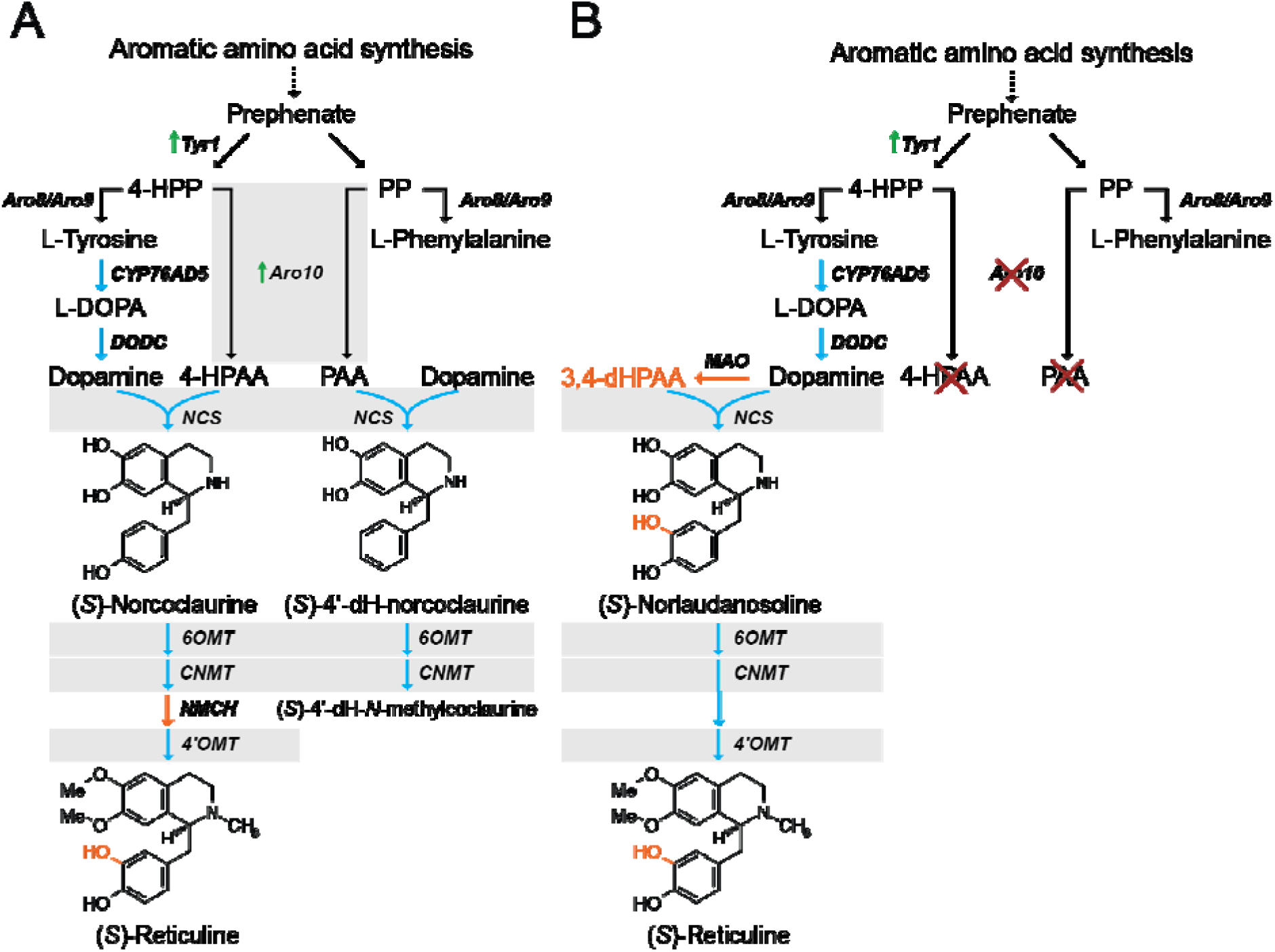
Two pathways for (*S*)-reticuline synthesis in yeast. **(A)** (*S*)-Reticuline synthesis in yeast proceeding through (*S*)-norcoclaurine. NCS catalyzes the condensation of dopamine and 4-HPAA to generate (*S*)-norcoclaurine, which is methylated and hydroxylated to form (*S*)-reticuline. Due to promiscuity of multiple pathway enzymes (grey boxes), (*S*)-4′-dehydronorcoclaurine and methylated derivatives are also produced when Aro10 is used to synthesize 4-HPAA. **(B)** (*S*)-Reticuline synthesis in yeast proceeding through (*S*)- norlaudanosoline, using largely the same enzymes as the (*S*)-norcoclaurine route. The key difference is the synthesis of the aldehyde 3,4-dHPAA from dopamine by MAO. The additional hydroxyl group on 3,4-dHPAA bypasses the requirement for NMCH-catalyzed hydroxylation. The two routes to (*S*)-reticuline’s 3′ hydroxyl group are highlighted in orange. Yeast native enzymes are indicated with black arrows; heterologous enzymes are indicated with blue arrows. Compound abbreviations: PEP, phosphoenolpyruvate; E4P, erythrose 4-phosphate; DHAP, 2-dehydro-3-deoxy-D-arabino-heptonoate 7-phosphate; 4-HPP, 4-hydroxyphenylpyruvate; 4-HPAA, 4-hydroxyphenylacetaldehyde; PP, phenylpyruvate, PAA, phenylacetaldehyde; 3,4-dHPAA, 3,4-dihydroxyphenylacetaldehyde. Enzyme abbreviations: DODC, L-DOPA decarboxylase; NCS, norcoclaurine synthase; MAO, monoamine oxidase; NMCH, *N*- methylcoclaurine hydroxylase; 6OMT, norcoclaurine 6-*O*-methyltransferase; CNMT, coclaurine *N*-methyltransferase; 4′OMT, 3′-hydroxyl-*N*-methylcoclaurine 4′ O-methyltransferase.

Synthesis of 4-HPAA from L-tyrosine through the intermediate tyramine circumvents the need for *ARO10* overexpression. Corresponding enzymes have been identified^14^ and expressed in yeast^15^, resulting in 0.08 g/L BIA synthesis in shake-flasks. An alternative, 4-HPAA-independent route to (*S*)-reticuline leverages 3,4-dihydroxyphenylacetaldehyde (3,4-dHPAA) to form (*S*)- norlaudanosoline. This pathway is largely identical to that of the (*S*)-norcoclaurine route; (*S*)- norlaudanosoline synthesis bypasses the need for a cytochrome P450-catalyzed hydroxylation reaction due to the presence of a second hydroxyl group on 3,4-dHPAA (Figure 1B, orange highlights). This has made norlaudanosoline the preferred route for BIA production in *E. coli*, in which several enzymes synthesizing 3,4-dHPAA from either L-DOPA^16^ or dopamine have been used. The highest titers (0.16 g/L reticuline) were achieved through the dopamine route, using monoamine oxidase from *Micrococcus luteus* (*Ml*MAO)^6^. Although *Ml*MAO was identified as a major bottleneck in *de novo* BIA synthesis in *E. coli*^6^, the success of the norlaudanosoline route for aldehyde synthesis highlights an opportunity to improve specificity and productivity of BIA production in yeast.

In this work, a yeast strain engineered to synthesize (*S*)-reticuline *via* (*S*)-norcoclaurine was retrofitted to synthesize (*S*)-reticuline *via* (*S*)-norlaudanosoline. 3,4-dHPAA synthesis from dopamine was achieved using human monoamine oxidase A (*Hs*MAO-A, hereafter MAO). The (*S*)-norlaudanosoline route to BIA synthesis in yeast enabled higher (*S*)-reticuline titers compared to the (*S*)-norcoclaurine route at a higher yield while almost eliminating undesirable condensation products. The heterologous pathway was further extended to the benzophenanthridines dihydrosanguinarine and sanguinarine, resulting in a titer of 653 mg/L in fed-batch fermentation surpassing previous microbially-produced titers 40-fold^15^.

## Results

### Additional copies of BIA modifying genes reduce accumulation of pathway intermediates

Previously, we reported *de novo* synthesis of benzylisoquinoline alkaloids (BIAs) in yeast at gram-per-liter scale, with the final strain (LP507) reaching 4.6 g/L (*S*)-reticuline in fed-batch fermentations^7^. However, mass spectrometry analyses revealed that, in addition to (*S*)-reticuline, strain LP507 also produced considerable quantities of other tetrahydroisoquinolines (THIQs). Based on total ion count, (*S*)-reticuline accounted for only 42% of all THIQs present in final fermentation broth (Figure 2A and 2B). An additional 25% was attributable to other benzyl THIQ BIA pathway intermediates upstream of (*S*)-reticuline: (*S*)-norcoclaurine, (*S*)-coclaurine, (*S*)-*N*-methylcoclaurine, and (*S*)-3**′**-hydroxy-*N*-methylcoclaurine (hereafter referred to as “other benzyl-THIQs”). The final 33% corresponded to THIQs resulting from the condensation of dopamine with other aldehydes (“non-BIA THIQs”).

**Figure 2.**
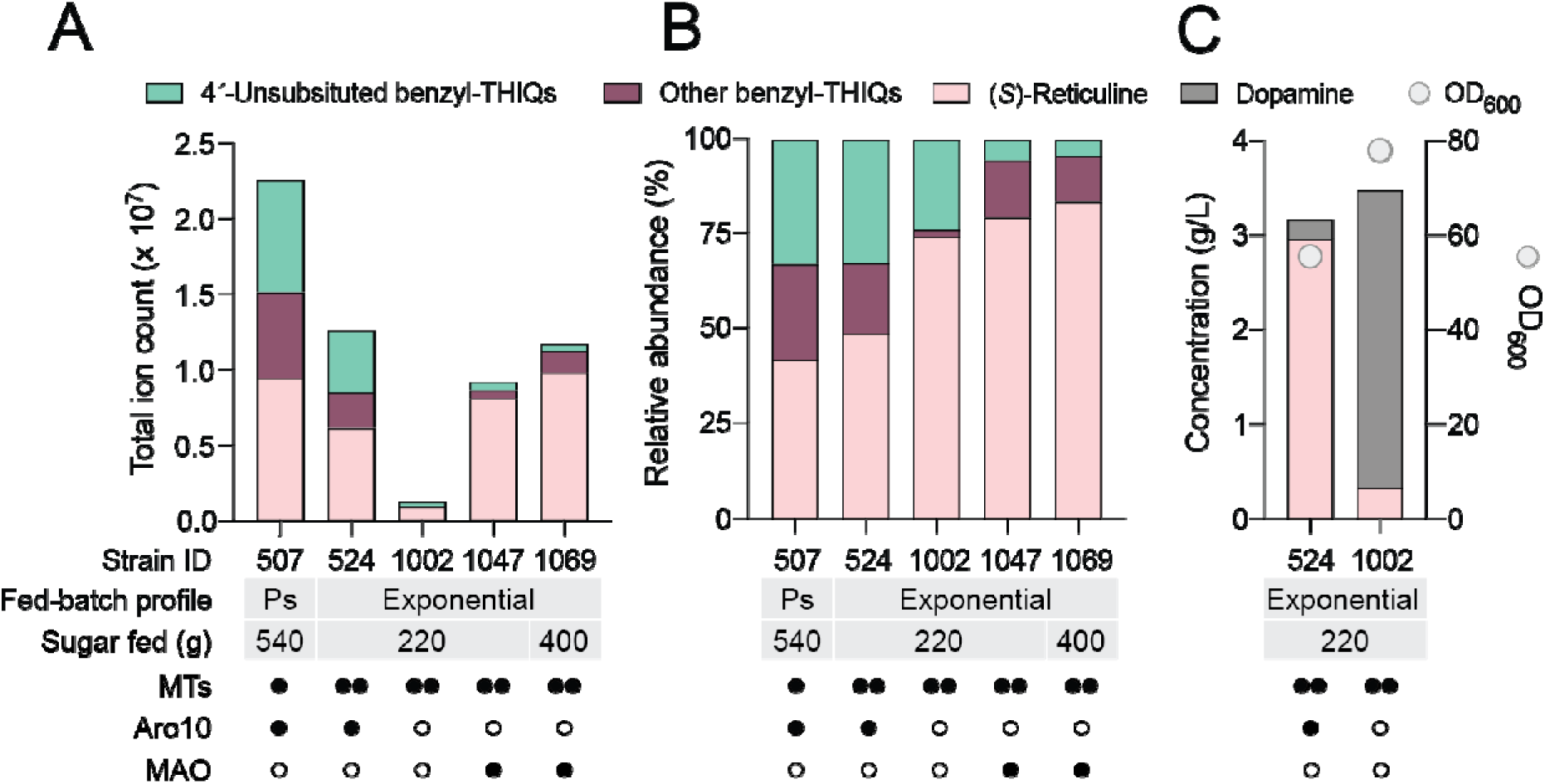
Effects of *ARO10* knockout in yeast synthesizing benzylisoquinoline alkaloids in fed-batch fermentation. **(A)** Absolute abundance of THIQs in select fed-batch fermentations. Sugar feeding was carried out either as a series of pulses controlled by off-gas analysis (Ps) or exponentially at a pre-set constant growth rate. End-point samples were analyzed for (*S*)-reticuline, benzyl-THIQ pathway intermediates, and off-pathway 4′-unsubstituted benzyl-THIQs originating from the condensation of dopamine and PAA. **(B)** Relative abundance of THIQs in the same fed-batch fermentations. **(C)** Impact of *ARO10* knockout on dopamine, (*S*)-reticuline, and biomass concentrations. The native and heterologous copies of *ARO10* were deleted from strain LP524 (strain LN1002) and both strains were grown in fed-batch fermentation with the same media and feeding profile. End-point samples were analyzed for OD_600_ and metabolite content. Symbols: ○ no enzyme; • one enzyme copy; •• two copies. Abbreviations: MTs, methyltransferases; MAO, monoamine oxidase.

To improve (*S*)-reticuline synthesis, we first sought to reduce the abundance of other benzyl-THIQs. Additional copies of pathway genes *6OMT*, *CNMT*, and *NMCH* were introduced to LP507, resulting in strain LP524. Analysis of total THIQs from fed-batch fermentation with strain LP524 revealed that relative abundance of other benzyl-THIQs fell from 25% to 18% (Figure 2B, strain LP524). However, non-BIA THIQs still comprised 33% of total peak area, indicating that further efforts should be focused on improving selectivity of THIQ synthesis.

### *ARO10* knockout minimizes *de novo* 4′-dehydroxynorcoclaurine synthesis

The dominant off-pathway side products detected in strain LP524 were compounds derived from the condensation of dopamine with phenylacetaldehyde (PAA): 4′- dehydroxynorcoclaurine (4**′**-dHN) and its methylated derivatives 4**′**-dehydroxycoclaurine and 4**′**- dehydroxy-*N*-methylcoclaurine (hereafter 4′-unsubstituted benzyl-THIQs; Figure 1A and Figure 2). Norcoclaurine synthase has a high degree of promiscuity for aldehydes, and will condense phenylacetaldehyde (PAA) with dopamine to produce 4**′**-dHN^12, 13^. Therefore, to limit 4′- unsubstituted benzyl-THIQ formation, the source of PAA synthesis in BIA-producing yeast should be identified and eliminated. Several 2-oxoacid decarboxylases have been identified in yeast^11^, with Aro10 among those contributing to PAA synthesis (Figure 1A)^17^. As *ARO10* was overexpressed in LP507 and LP524, we hypothesized that this enzyme was responsible for high levels of PAA production in our strains.

To test this hypothesis, both the native and overexpressed copies of *ARO10* were deleted in strain LP524 to generate strain LN1002. Strains LP524 and LN1002 were both grown in fed batch fermentations using the same feeding profile and amount of sugar. Under these conditions, LP524 synthesized 3.0 g/L of (*S*)-reticuline and 0.2 g/L dopamine, whereas the *aro10*Δ LN1002 synthesized 3.1 g/L of dopamine and 0.33 g/L (*S*)-reticuline (Figure 2C). The 90% reduction in (*S*)-reticuline levels demonstrates the dominant contribution of Aro10 on 4-HPAA synthesis in the parent strain. Importantly, mass spectrometry revealed almost complete ablation of 4′- unsubstituted benzyl-THIQs (Figure 2A and Figure 2B), confirming that Aro10 was also largely responsible for PAA synthesis in LP524. Eliminating Aro10 activity also improved biomass accumulation in the bioreactor; strain LN1002 grew to a final OD_600_ of 78 compared to 55 for strain LP524.

### Expression of *MAO* in an *ARO10* knockout strain restores BIA synthesis

With high dopamine titers and low off-target metabolite accumulation, the *aro10*Δ LN1002 provided a clean starting point for studying an alternative route to aldehyde generation for BIA synthesis. We opted to explore the production of 3,4-dihydroxyphenylacetaldehyde (3,4-dHPAA), which can be synthesized from and subsequently condensed with dopamine to form the norcoclaurine analog norlaudanosoline (Figure 1B). We chose to examine human monoamine oxidase A (MAO), which was previously suggested as a candidate for (*S*)-norlaudanosoline synthesis in yeast^18^.

First, we determined the impact of promoter strength of MAO on (*S*)-reticuline synthesis and strain fitness in microtiter plate format. *MAO* codon-optimized for expression in yeast was introduced to LN1002 under four well-characterized promoters spanning several orders of magnitude of relative strength^19^. MAO activity was assessed by growing strains in deep-well plates and measuring their metabolite profile. *MAO* expression reduced dopamine titers in LN1002 in a promoter strength-dependent manner (Figure 3A). Hydroxytyrosol, which is produced when 3,4-dHPAA is reduced by yeast oxidoreductases (Figure 4A), increased with promoter strength, indicating that MAO was successfully generating 3,4-dHPAA. Crucially, successful restoration of (*S*)-reticuline synthesis in a promoter-dependent manner also confirmed that 3,4-dHPAA can not only be produced but also condensed with dopamine in yeast to produce (*S*)-norlaudanosoline, which can then be methylated to form (*S*)-reticuline.

**Figure 3.**
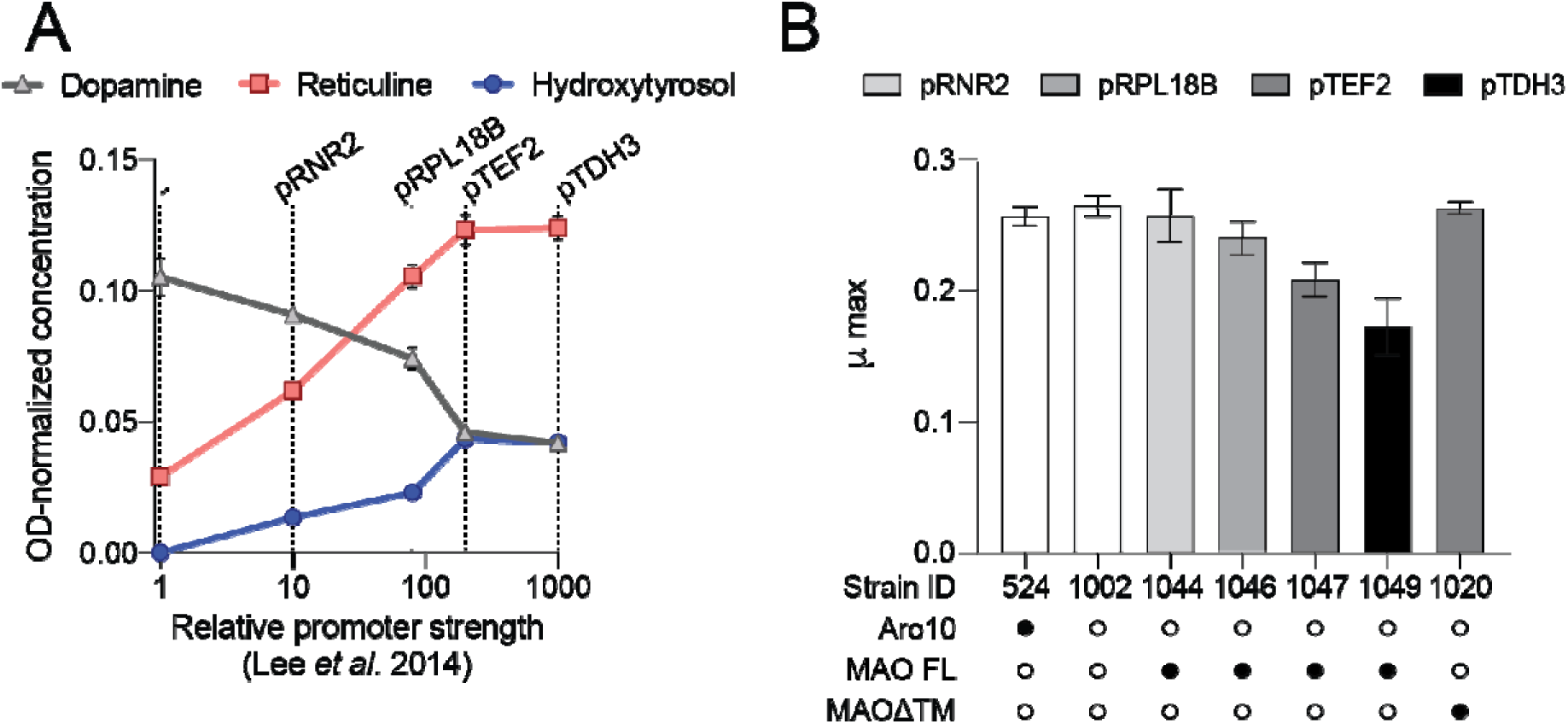
Promoter titration of *MAO* in an *ARO10* knockout background. Strain LN1002 lacking Aro10 was used to express *MAO* under the control of *pRNR2*, *pRPL18B*, *pTEF2* or *pTDH3*. **(A)** Cultures were grown in rich media in deep well plates and then analyzed for metabolite profile and final OD_600_. Metabolite profile of strains presented as g/L normalized to OD_600_. Error bars refer to mean and standard deviation of n=2 biological replicates. **(B)** Cultures were grown in minimal media in microtiter plates in a plate reader, and maximum specific growth rate was obtained from the growth curves. Promoter strength driving MAO expression is indicated by grey shade. Abbreviations: MAO FL, full length MAO; MAOΔTM, MAO with transmembrane helix deleted. Symbols: ○ no enzyme; • one enzyme copy. Error bar refer to mean and standard deviation of n=3 biological replicates.

**Figure 4.**
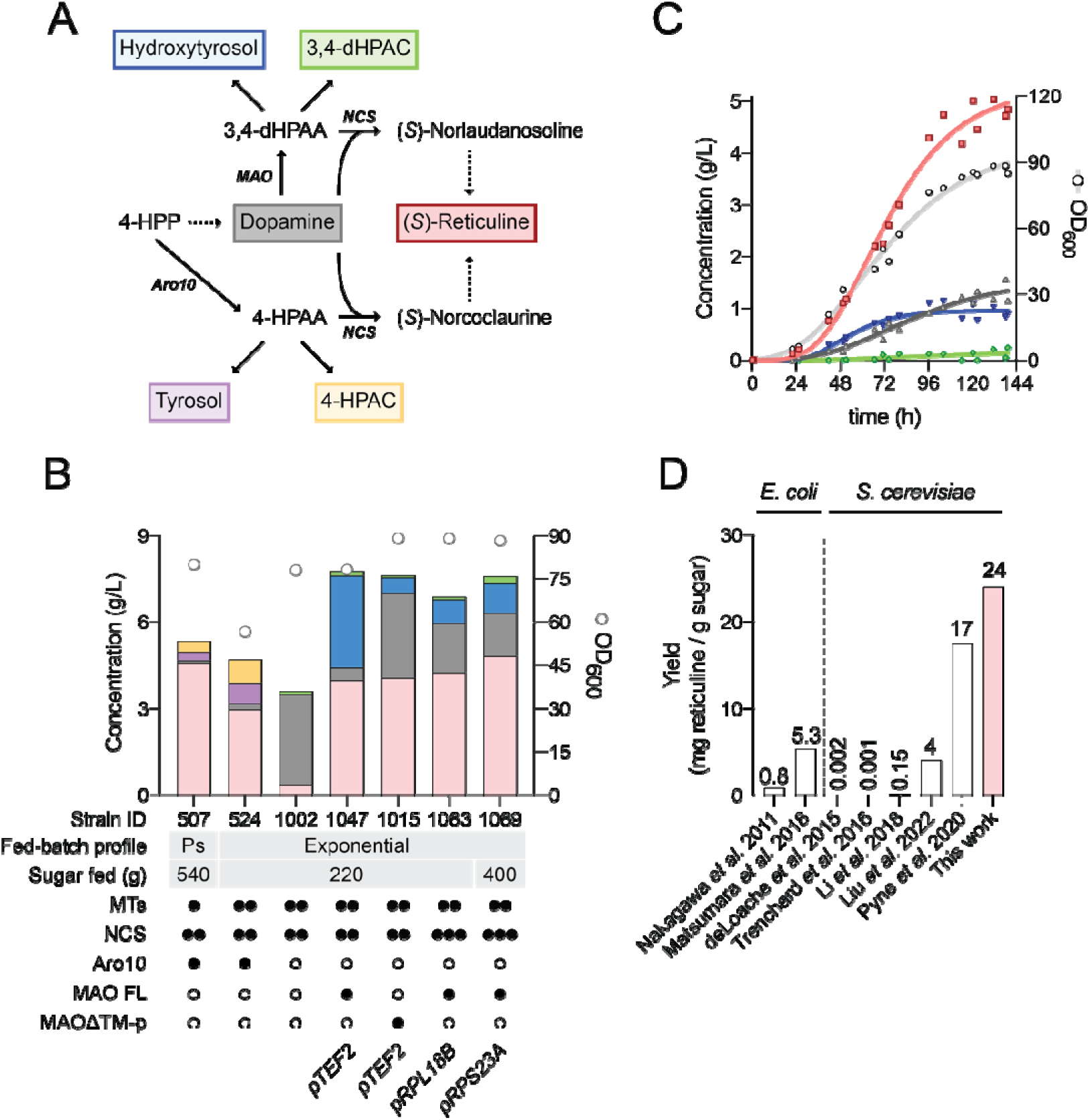
Optimization of (*S*)-reticuline production in fed-batch fermentation. **(A)** Pathways for fusel acid and alcohol production in strains synthesizing 4-HPAA and 3,4-dHPAA. Dashed arrows refer to multiple enzymatic reactions. **(B)** End-point analysis of quantifiable metabolites of BIA-producing strains grown in fed-batch fermentation. Strains were grown in batch phase until sugar was exhausted, then one of two sugar feeding fed-batch profiles was used pulsed (Ps) or exponential. Pulsed fed-batch media contained 500 g sucrose/L, exponential fed-batch media contained 360 g sucrose/L. Bar graph colors correspond to metabolites in Panel A. **(C)** Time course of fed-batch fermentation with LN1069, with *MAO* expression under control of *pRPS23A*. Two replicates are plotted with a best fit curve included to guide the eye. Line graph colors correspond to metabolites in Panel A. **(D)** *De novo* reticuline yields reported in the literature expressed as mg reticuline/g sugar. Compound abbreviations: 4-HPP, 4-hydroxyphenylpyruvate; 4-HPAC, 4-hydroxyphenylacetate; 3,4-dHPAA, 3,4-dihydroxyphenylacetaldehyde; 3,4-dHPAC, 3,4-dihydroxyphenylacetate. Enzyme abbreviations: MTs, methyltransferases; NCS, norcoclaurine synthase. Symbols: ○ no enzyme; • one enzyme copy; •• two copies; ••• three copies.

Growth rate assays performed in parallel with metabolite analyses revealed that strains’ maximum specific growth rate (µ_max_) decreased with increased *MAO* expression (Figure 3B). In humans, MAO harbors a 30 amino acid C-terminal transmembrane helix anchored in the outer mitochondrial membrane^20, 21^. We investigated whether mitochondrial localization of MAO affected growth rate by removing the transmembrane helix and expressing this variant (MAOΔTM) in yeast under control of the *TEF2* promoter (strain LN1020). Removal of transmembrane helix restored µ_max_ to the level observed in strains not expressing MAO (Figure 3B), but it did not impact MAO localization (Supplemental Figure 1). We also observed that the activity of MAOΔTM was lower than its full-length counterpart (Supplemental Figure 2A). Taken together, we conclude that the improved growth rate of cells expressing MAOΔTM was due to reduced enzymatic activity and not due to altered localization of the enzyme. This corroborates a report from 1994 in which MAOΔTM was demonstrated to maintain localization at the mitochondria while having difficulty forming disulfide bridges essential for full enzymatic activity^22^.

Synthesis of 3,4-dHPAA from dopamine also produces a molar equivalent of hydrogen peroxide, which may impose a burden on the cell and reduce µ_max_. To mitigate these effects, we attempted to target MAO to the peroxisome using the peroxisomal targeting tag ePTS1 (ref ^23^) (LN1015; MAOΔTM-p). However, there was no evidence that MAO was localized to the peroxisome, with or without removal of the transmembrane helix (Supplemental Figure 1 and 3). Further studies are required to better understand how to manipulate MAO localization without also perturbing enzyme activity.

### *MAO* driven by *pTEF2* limits off-target THIQ synthesis in fed-batch fermentation

A key challenge of BIA production in *S. cerevisiae* is its propensity to transform precursor aldehydes into fusel alcohols or acids^7^. Aldehydes may be oxidized or reduced by oxidoreductases depending on the redox environment of the cell^17^. In our prior study, significant residual 4-HPAA catabolic activity remained in strain LP507, even with the deletion of seven oxidoreductases. Oxidation of 4-HPAA into 4-hydroxyphenylacetic acid (4-HPAC) was especially persistent. In that study we circumvented this issue by utilizing a fed-batch protocol that promoted periodic production and reuptake of ethanol a series of sugar pulses controlled through off-gas analysis intended to maintain an environment where yeast has limited capacity to oxidize 4-HPAA^24^. A downside of this protocol is carbon loss in the repeated phases of fermentative growth and ethanol production. Moreover, utilizing a simple, sugar-limited feeding profile would be a preferred solution for BIA synthesis in industrial settings. Thus, we sought to establish an exponential fed-batch regime in *MAO*-expressing strains, using the experiment with strain LP524 as a baseline for comparison.

Compared to LP524, strain LN1047 expressing *MAO* from the *TEF2* promoter produced more (*S*)-reticuline (3.0 vs 4.0 g/L; Figure 4B). Crucially, strain LN1047 made fewer 4′-unsubstituted benzyl-THIQs (5% vs 33% of total peak area; Figure 2B). In total, (*S*)-reticuline comprised 80% of THIQs by peak area, compared to the 49% of LP524. These initial results demonstrated that BIA synthesis through 3,4-dHPAA successfully reduced production of off target condensation products. However, the presence of 3.2 g/L hydroxytyrosol, 0.16 g/L 3,4-dHPAC, and 0.44 g/L dopamine indicated a need for additional strain optimization.

### Balancing expression of pathway branch point enzymes enhances norlaudanosoline synthesis

Rerouting (*S*)-reticuline synthesis using MAO improved selectivity of THIQ synthesis (Figure 3A and 3B), but the accumulation of dopamine and the 3,4-dHPAA-derived hydroxytyrosol posed a new problem (Figure 4B). A delicate balance must be achieved in the synthesis of dopamine, 3,4-dHPAA, and (*S*)-norlaudanosoline (Figure 4A): MAO activity for the conversion of dopamine to 3,4-dHPAA must not be too high or too low, while NCS activity must be sufficient to condense dopamine and 3,4-dHPAA before the former exits the cell or scavenging oxidoreductases convert the latter to hydroxytyrosol/3,4-dHPAC. When *MAO* was expressed from the *TEF2* promoter, the concentration of hydroxytyrosol was 6-fold higher than dopamine in fed-batch fermentation, suggesting excessive MAO activity. Thus, two options were explored to alter the ratio of dopamine and 3,4-dHPAA: using the MAO variant MAOΔTM-p, a variant with reduced activity compared to wild-type MAO and varying the strength of the promoter driving *MAO* expression.

In fed-batch fermentation, *MAOΔTM-p* under control of the *TEF2* promoter (strain LN1015) resulted in the synthesis of 4 g/L (*S*)-reticuline, in this instance with the additional accumulation of 3 g/L dopamine and 0.6 g/L hydroxytyrosol (Figure 4B), pointing to insufficient activity of MAO. Additionally, the continued accumulation of both dopamine and hydroxytyrosol in fermentations with strains LN1047 and LN1015 suggested that NCS activity may not be sufficient for efficient condensation of dopamine and 3,4-dHPAA (Figure 4B). *NCS* expression negatively impacts strain fitness, which can be alleviated by peroxisomal sequestration^23^. Thus, an extra copy of peroxisomally-targeted NCS was introduced to strains moving forward.

To address precursor imbalance, we probed weaker promoters driving full-length *MAO* expression. With *MAO* under the control of the *RPL18B* promoter, strain LN1063 produced 4.25 g/L (*S*)-reticuline while continuing to produce 1.5 g/L dopamine and 0.5 g/L hydroxytyrosol (Figure 4B). Although this resulted in a slight improvement in (*S*)-reticuline yield, dopamine was still 3 times as abundant as hydroxytyrosol, indicating that optimal MAO activity in this strain may require a promoter with strength between *pRPL18B* and *pTEF2*.

The yeast MoClo collection of characterized promoters does not include any with strength intermediate to *pRPL18B* and *pTEF2* (ref ^19^). A 2010 report from Canelas *et al*. contains RNASeq data of two common yeast strains, CEN.PK and S288C, grown in both shake-flask and sugar-limited chemostat conditions^25^. We searched these data for promoters whose strength was between those of *pRPL18B* and *pTEF2* in both CEN.PK and S288C (the strains in this study are based on the latter) in both shake-flask and sugar-limited conditions. Three promoters, *pRPL39*, *pHTB1*, and *pRPS23A*, were selected. We confirmed that these promoters were indeed intermediate to *pRPL18B* and *pTEF2*, as determined by dopamine, hydroxytyrosol, and (*S*)- reticuline abundance in 96-well plate format (Supplemental Figure 2B). From this set, strain LN1069, expressing full length *MAO* from *pRPS23A* as well as a second copy of peroxisomally targeted *NCS*, was selected for further characterization in a 3L bioreactor (Figure 4C).

In fed-batch cultivation, strain LN1069 synthesized 4.8 g/L (*S*)-reticuline, with an additional 1.5 g/L dopamine and 1.0 g/L hydroxytyrosol (Figure 4B and 4C). In this experiment, we increased total sucrose fed to 400 g, corresponding to an overall yield of 24 mg (*S*)- reticuline/g sucrose (Figure 4D). By mass spectrometry, (*S*)-reticuline comprised 83% of THIQs with another 12% attributable to benzyl-THIQs and 5% to 4′-unsubstituted benzyl-THIQs (Figure 2B).

### *De novo* synthesis of dihydrosanguinarine *via* (*S*)-norlaudanosoline

Previously, we introduced a pathway into yeast to convert supplemented (*S*)- norlaudanosoline to dihydrosanguinarine^26^. In that work, we identified a pathway bottleneck between (*S*)-scoulerine and (*S*)-stylopine synthesis, which was resolved in a follow-up report^27^. In the present work, all six enzymes were introduced to a yeast strain synthesizing (*S*)-reticuline *de novo* via (*S*)-norlaudanosoline (strain LN1015) (Figure 5A).

**Figure 5.**
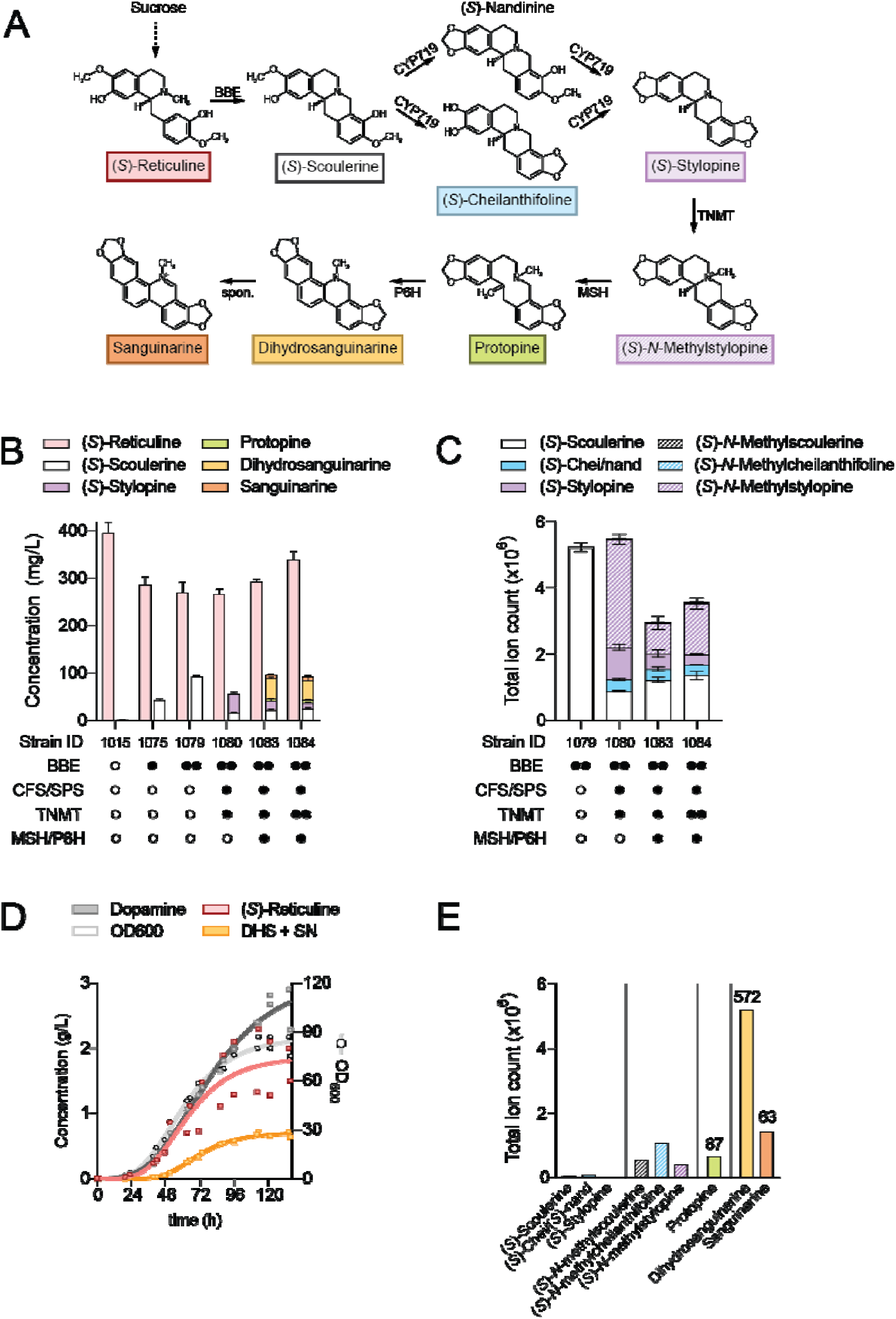
*De novo* dihydrosanguinarine synthesis in yeast. **(A)** Biosynthesis of dihydrosanguinarine from (*S*)-reticuline. **(B)** Stepwise construction of a *de novo* dihydrosanguinarine strain. Pathway enzymes were integrated into LN1015, a strain synthesizing (*S*)-reticuline *via* (*S*)-norlaudanosoline. Metabolites were extracted and, where possible, quantified by LC-MS. Error bars refer to mean and standard deviation of n=3 biological replicates. **(C)** Metabolite profile of intermediates between (*S*)-scoulerine and (*S*)-*N*- methylstylopine. Error bars refer to mean and standard deviation of n=3 biological replicates. **(D)** Fed-batch fermentation of LN1084, a strain synthesizing dihydrosanguinarine *de novo*. Samples were regularly collected for OD_600_ and metabolite analysis. Two replicates are plotted with a best fit line to guide the eye. **(E)** Metabolite profile of target and intermediate compounds at the final time point of a representative fed-batch fermentation in panel (D). Concentrations of quantifiable metabolites are indicated in mg/L. Abbreviations: BBE, berberine bridge enzyme; CFS, (*S*)- cheilanthifoline synthase; SPS, (*S*)-stylopine synthase; TNMT, (*S*)-tetrahydroprotoberberine *N*- methyltransferase; MSH, (*S*)-*N*-methylstylopine hydroxylase; P6H, protopine 6-hydroxylase; DHS, dihydrosanguinarine; SN, sanguinarine. Symbols: ○ no enzyme; • one enzyme copy; •• two copies.

Introduction of one copy of the gene encoding the berberine bridge enzyme *BBE* to strain LN1015 resulted in 15% conversion of (*S*)-reticuline to (*S*)-scoulerine in deep-well plates (Figure 5B, strain LN1075). A second copy of *BBE* improved conversion to 25% (LN1079). The next three pathway enzymes, (*S*)-cheilanthifoline synthase (CFS), (*S*)-stylopine synthase (SPS) and (*S*)-tetrahydroprotoberberine *N*-methyltransferase (TNMT) were introduced into strain LN1079, generating strain LN1080 (Figure 5B). In our 2016 report, these three enzymes converted 100% of supplemented (*S*)-scoulerine to (*S*)-*N*-methylstylopine. Here, conversion was incomplete – 15 mg/L of residual (*S*)-scoulerine was detected, along with some residual (*S*)-cheilanthifoline. A minor bottleneck at (*S*)-stylopine methylation was also observed. Finally, the two cytochromes P450 MSH and P6H were introduced to LN1080, generating a strain of yeast producing dihydrosanguinarine *de novo* from sucrose (Figure 5B, LN1083). In deep-well plates, LN1083 synthesized 40 mg/L dihydrosanguinarine. Additionally, 7 mg/L sanguinarine was produced, presumably through spontaneous oxidation of dihydrosanguinarine. The impact of a second copy of *TNMT* was also probed (strain LN1084). While there was no improvement in dihydrosanguinarine titers, we observed a decrease in (*S*)-stylopine and an increase in (*S*)-*N*- methylstylopine, indicating that the extra copy of *TNMT* was effective (Figure 5C).

Growth of strain LN1084 in sugar-limited fed-batch conditions resulted in the synthesis of 572 mg/L dihydrosanguinarine and an additional 63 mg/L sanguinarine, for a combined output of 635 mg/L (Figure 5D). Additionally, 2.2 g/L (*S*)-reticuline and 87 mg/L protopine accumulated in the broth. BBE remained the rate-limiting step in flux through the dihydrosanguinarine pathway in fed-batch conditions, in accordance to experiments in deep-well plates. (*S*)-Scoulerine, (*S*)-cheilanthifoline, and (*S*)-stylopine were almost undetectable, whereas the three *N*-methylated equivalents accumulated (Figure 5E).

### *De novo* synthesis of side products upon introduction of dihydrosanguinarine synthesis

Several novel peaks were observed upon co-expression of enzymes to convert (*S*)- reticuline to dihydrosanguinarine. Two distinct peaks with exact [M+H]^+^ 356.1490 (C_20_H_21_NO_5_) with similar retention times were identified (Figure 6A). The chemical formula and MS^2^ profile suggested that these compounds are closely related to protopine hunnemanine and izmirine, (Figure 6B). These compounds are proposed to be derived from the activity of MSH on (*S*)-(*N*)- methylcheilanthifoline and (*S*)-(*N*)-methylnandinine rather than (*S*)-(*N*)-methylstylopine (Figure 6C). Assuming equal ionization efficiency to protopine, these peaks could represent 30 mg/L of lost carbon.

**Figure 6.**
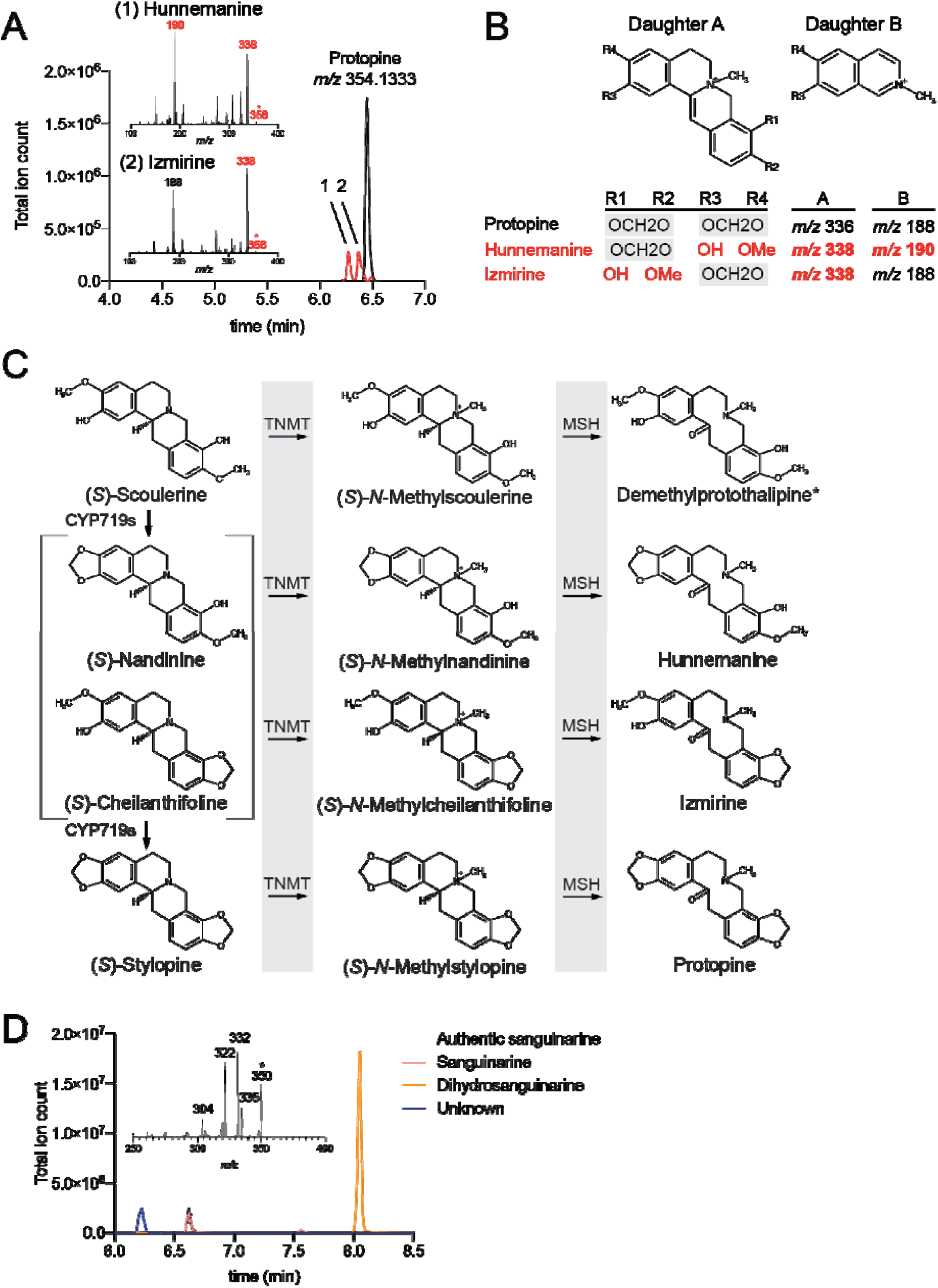
Unidentified metabolites in dihydrosanguinarine fermentation samples of strain LN1084. Broth from the fed-batch fermentation of strain LN1084, synthesizing *de novo* dihydrosanguinarine, was extracted and analyzed by HPLC-FT-MS/MS. **(A)** Two unknown peaks with exact *m/z* [M+H]^+^ of 356.1490 were observed; chromatograms and MS^2^ fragmentation patterns are displayed, with MS^2^ daughter ions differing by 2 *m/z* from protopine highlighted in red. **(B)** Two MS^2^ fragmentation ions characteristic to protopines, which can be used to distinguish unknowns 1 and 2. **(C)** Synthesis of hunnemanine and izmirine is proposed to proceed through activity of the enzyme MSH on the *N*-methylated pathway side-products *N*- methylnandinine and *N*-methylcheilanthifoline, respectively. **(D)** A peak with exact *m/z* [M+H]^+^ of 350.1025 was observed; chromatograms and MS^2^ fragmentation patterns are displayed.

We detected an additional unknown compound with exact [M+H]^+^ of 350.1025 (C_20_H_17_NO_5_) (Figure 6D). The compound is likely a benzophenanthridine like sanguinarine, or a dihydrobenzophenanthridine like dihydrosanguinarine, displaying characteristic losses of [M-Me]^+^ (15 *m*/*z*) and [product ion+H-CO]^+^ (28 *m/z*) as well as a fragment corresponding to the exact mass of sanguinarine (*m/z* 322)^28^. Its chemical formula corresponds to a hydroxydihydrosanguinarine. However, its MS^2^ profile does not match that of 10-hydroxydihydrosanguinarine^29^. Alternatively, 6-hydroxydihydrosanguinarine (also known as sanguinarine pseudobase) is a naturally-occurring form of sanguinarine that exists in equilibrium with sanguinarine and sanguinarine dimer at physiological pH^30^. An authentic MS^2^ profile of sanguinarine pseudobase could not be obtained for validation of this hypothesis. Sanguinarine pseudobase is not observed in authentic sanguinarine standard (Figure 6C), but the standard was not held at physiological pH. Assuming ionization efficiency equal to that of sanguinarine, this compound could represent an additional 100 mg/L of material derived from (*S*)-reticuline either on-pathway if it is sanguinarine pseudobase or off-pathway if it is dihydrosanguinarine hydroxylated at another position.

## Discussion

In 2020, we reported the first gram-per-liter-scale synthesis of the benzylisoquinoline alkaloid (BIA) (*S*)-reticuline in a microbial host^7^. While a major improvement over earlier results, the pathway suffered from excessive accumulation of side-products formed through the concerted activity of the Ehrlich pathway enzyme Aro10 and norcoclaurine synthase (NCS). Here, we employ an alternative pathway by rerouting (*S*)-reticuline synthesis through (*S*)- norlaudanosoline using the enzyme monoamine oxidase A from humans (MAO, Figure 1B). This MAO-enabled route to (*S*)-reticuline synthesis results in improved yields (24 vs 17 mg/g glucose) and higher titers (4.8 vs 4.6 g/L) while dramatically reducing off-target condensation products: 5% vs 33% in comparison to previous report.

Enzyme promiscuity was a frequent theme throughout this work. Yeast could catabolize the non-native aldehyde 3,4-dHPAA. 4′-unsubstituted THIQ synthesis required concerted activity of Aro10, NCS, and the methyltransferases 6OMT and CNMT were capable of further modifying this off-pathway THIQ (Figure 1). Further, upon introduction of *de novo* dihydrosanguinarine synthesis, multiple side products including hunnemanine and izmirine were discovered, which required promiscuity of both TNMT and MSH (Figure 5E). The synthesis of 4′-unsubstituted THIQs was largely resolved in this work, but additional challenges remain.

One general strategy for addressing enzyme promiscuity is to ensure that the heterologous pathway of interest is functioning with as linearly and with minimal bottlenecks. This is traditionally achieved using standard approaches such as copy number titration and/or identification of orthologous enzymes with superior specificity and/or activity in yeast. Heterologous enzymes may need modifications to function well in a yeast host. For example, the NCS variant used here was previously improved through deletion of an N-terminal signal peptide^7^. Other options to limit the impact of promiscuity include enzyme co-localization through scaffolding, fusion, or compartmentalization into natural or synthetic organelles^31, 32^.

A complementary strategy is to ensure that the native metabolism of the yeast host is fine-tuned to be as compatible with the heterologous pathway as possible. For instance, the methyltransferases converting (*S*)-norlaudanosoline to (*S*)-reticuline depend on *S*- adenosylmethionine (SAM). It may be beneficial to introduce SAM cofactor engineering^33^ to facilitate conversion of (*S*)-norlaudanosoline to the triple-methylated (*S*)-reticuline. This could reduce the accumulation of benzyl-THIQ pathway intermediates, which comprised 12% of total THIQs in the final strain LN1069 (Figure 2B).

Paradoxically, we successfully took advantage of enzyme promiscuity to improve pathway specificity; rerouting (*S*)-reticuline synthesis through 3,4-dHPAA required concerted promiscuity of NCS, 6OMT, CNMT and 4′OMT to succeed (Figure 1B). By virtue of this, strain LN1069 produced just 5% 4′-unsubstituted benzyl-THIQs, down from 33% in LP524 (Figure 2B). However, strains producing (*S*)-reticuline through (*S*)-norcoclaurine generated relatively little 4-HPAA-derived fusel alcohols and acids, while in strains leveraging the (*S*)-norlaudanosoline route accumulation of fusel products remained a major issue (Figure 2B, Figure 4A and 4B). This indicates that problems with off-target THIQ and precursor degradation were effectively inverted in the two lineages. Somewhat surprisingly, the major degradation product of 3,4-dHPAA in strain LN1069 was hydroxytyrosol, a non-native fusel alcohol (Figure 4B), in spite of deletion of 7 oxidoreductases which dramatically reduced accumulation of the closely related tyrosol in LP524 (reference 7). This illustrates the redundancy and efficacy of yeast’s Ehrlich pathway^17^. Additional deletions may be required to ensure efficient function of this key branch point, although some enzymes demonstrated to act on aromatic aldehydes, such as Ald4p (reference 7), cannot be easily removed from an industrially-viable host.

Strains expressing *MAO* also accumulated more unreacted dopamine than strains over expressing *ARO10* in fed-batch fermentations (Figure 4B). Recapturing the carbon lost to fusel products and dopamine could result in as much as a 50% increase in (*S*)-reticuline titer and productivity metrics in these strains. The pathway branch point at dopamine, 3,4-dHPAA, and (*S*)-norlaudanosoline (Figure 4A) must be approached delicately. Although increasing NCS expression through additional gene copy number increase may prove useful to decrease precursor accumulation, the burden of high NCS load may limit the utility of this approach^23^. Alternatively, entry into the aromatic amino acid pathway could be dynamically controlled using a transcriptional biosensor specific to a pathway side product such as hydroxytyrosol^34^. We also noticed dopamine concentration continued to rise towards the end of fermentation while those of hydroxytyrosol and (*S*)-reticuline plateaued (Figure 4C). This could be attributed to decreasing MAO activity or changes in expression levels of other native and non-native enzymes. A broader scan of promoters for *MAO* expression may identify a candidate that is more appropriate for maintaining consistent BIA production independently of growth rate^25, 35^.

Biomass yield began to plateau towards the end of fed-batch fermentations of BIA synthesizing strains, whether they produce (*S*)-reticuline via 3,4-HPAA (Figure 4C) or 4-HPAA (Supplemental Figure 4). As growth plateaued so did (*S*)-reticuline synthesis, suggesting that BIA production is at least partially growth-coupled. The cause of growth inhibition should be identified and addressed, as it represents a major obstacle to further productivity improvements. It is likely that the heavily engineered strain background has multiple sources of stress, the effects of which are exacerbated over the course of a fermentation. LP524, producing (*S*)- reticuline through 4-HPAA, reached a lower biomass density than the *aro10*Δ LN1002; the final OD stayed higher even when MAO was introduced and (*S*)-reticuline synthesis was restored (Figure 4B). This suggests some degree of stress from *ARO10* expression in a manner independent of BIA synthesis, such as the presence of 4′-unsubstituted THIQs or excess aldehyde produced through Aro10 in an oxidoreductase knockout background. Higher *MAO* expression level was accompanied with a reduction of maximum specific growth rate in 96-well plates. Redirection of MAO to the peroxisome was unsuccessful (Supplemental Figures 1 and 3), but additional approaches to mitigating hydrogen peroxide stress, such as catalase overexpression^36^ could be explored. Deleterious effects of elevated intracellular BIAs cannot be ruled out, in which case engineering BIA export may alleviate some growth inhibition^37^.

The BIA sanguinarine is a primary active ingredient in Sangrovit®, a probiotic used in animal husbandry. To demonstrate the commercial potential of our yeast BIA platform, an (*S*)- reticuline producing strain (LN1015) was modified to produce dihydrosanguinarine *de novo* from sugar. In fed-batch fermentation, we produced 635 mg/L of dihydrosanguinarine and its oxidized derivative sanguinarine from sucrose. This represents the highest titer of those BIAs produced in microbes by a factor of 40 (reference 15). It was also a sufficient quantity to serve as a probiotic for approximately one ton of fish feed^38, 39^. Here we also describe, to the best of our knowledge, the first *de novo* synthesis of the protopines hunnemanine and izmirine. These compounds were synthesized due to a minor bottleneck at (*S*)-stylopine synthesis. While this bottleneck was previously resolved, it may have reappeared due to the increased scale of *de novo* (*S*)-scoulerine synthesis herein; over 2 mM of (*S*)-scoulerine-derived BIAs were produced using strain LN1084 in fed-batch fermentation (Figure 5E) vs 5 μM (*S*)-scoulerine supplementation in previous work. Conversion of (*S*)-reticuline to (*S*)-scoulerine was also incomplete, highlighting a previously-described key bottleneck at BBE^15^. BBE has been shown to be poorly soluble in *E. coli*. This was ameliorated by generating an N-terminal maltose-binding fusion protein, resulting in an 80-fold improvement in BBE activity *in vivo*^40^. This fusion protein strategy may also prove beneficial in our strains, potentially enabling commercially-viable production of (*S*)-scoulerine derived BIAs including protoberberines, protopines, benzophenanthridines, and phthalideisoquinolines in the future.

In summary, we engineered yeast to synthesize (*S*)-reticuline *via* (*S*)-norlaudanosoline, enabling improved productivity metrics compared to the previously reported (*S*)-norcoclaurine route. We also extended the heterologous pathway to dihydrosanguinarine, resulting in highest titer of this commercial BIA in a microbial host. We highlight the importance of monitoring of both of on and off-pathway metabolites, challenges with reducing catabolism of pathway intermediates, and balancing enzyme expression levels in branch points in THIQ biosynthesis in yeast.

## Acknowledgments

We thank Marcos DiFalco for assistance with maintenance of mass spectrometers. This study was financially supported by an NSERC-Industrial Biocatalysis Network (IBN) grant and an NSERC Discovery grant. L.N. was supported by a FRQNT DE Doctoral Research Scholarship for Foreign Students, M.E.P. was supported by an NSERC Postdoctoral Fellowship, K.K. was supported by a Concordia University Horizon Postdoctoral Fellowship. K.-H.S. and J.E.D. were supported by NSF CBE 2104261 and NSF DBI 1548297. V.J.J.M. is supported by a Concordia University Research Chair.

## Author contributions

L.N., K.K., M.E.P., K.H.S., J.E.D. and V.J.J.M. designed the research. L.N., M.E.P., and K.H.S. performed the experiments. K.K. assisted in fermentation analysis. J.E.D and V.J.J.M supervised the research. L.N., K.K., K.H.S. and V.J.J.M wrote the manuscript with editing help from M.E.P. and J.E.D.

## Competing financial interests

The authors declare no competing financial interests.

## Additional information

Supplementary information is available in the online version of the paper. Correspondence and requests for materials should be addressed to V.J.J.M.

## Data availability

The data that support the findings of this study are available from the authors upon reasonable request.

## Materials & Methods

### Yeast and *E. coli* growth conditions

*E. coli* was grown in liquid Luria Broth (10 g/L peptone, 5 g/L yeast extract, 10 g/L sodium chloride, LB; Fisher Bioreagents) at 37°C with shaking at 200 rpm. *E. coli* transformations were selected on solid LB with 2% agar with antibiotics supplied as necessary.

For genetic manipulation, yeast was grown in liquid yeast peptone dextrose (20 g/L peptone, 20 g/L dextrose, 10 g/L yeast extract, YPD; Sigma Aldrich) at 30°C with shaking at 200 rpm. Yeast transformations with Cas9-containing plasmids were selected on solid YPD with 2% agar, 200 µg/mL G418, and 200 µg/mL hygromycin. For assessment of BIA synthesis in 96-well plate format, yeast was grown in 2x synthetic complete media (2x SC: 13.6 g/L Difco Yeast Nitrogen Base (YNB), 3.84 g/L yeast synthetic drop-out medium supplements without histidine (Millipore-Sigma), 152 mg/L histidine, 40 g/L sucrose) with shaking at 400 rpm in 96-well 2 mL deep-well plates overnight, followed by a 1:50 back dilution into fresh 2x SC for 3 days. For assessment of yeast growth in 96-well plate format, strains were grown in 2x SC overnight with shaking at 400 rpm, followed by a 1:100 back dilution into YNB with 20 g/L sucrose supplemented with 76 mg/L methionine and 76 mg/L histidine. Prior to growth in fermenter, strains were transformed with a plasmid complementing methionine and histidine auxotrophies and selected on solid 1x SC media lacking histidine (SC-His).

### Strain construction

Gene knockouts and genomic integrations were introduced to yeast *via* CRISPR-directed homologous recombination. A plasmid harboring Cas9 and an empty guide RNA transcription cassette was linearized by *Not*I/*Bsa*I double digestion (New England Biolabs) and transformed into yeast together with a linear piece of DNA containing the guide RNA sequence flanked on either side by homology to the plasmid, which resulted in *in vivo* plasmid assembly. Linear DNA containing the guide was generated by PCR. Guide RNAs for gene knockout were selected based on a combined score from the online tools Yeast CRISPRi^41^ and CCTOP^42^. Gene knockouts were generated through co-transformation of a linear fragment containing 40 bp homology to either side of the gene of interest. Gene integrations were targeted to genomic regions previously identified to promote high-level *GFP* expression^43, 44^. Gene integrations were introduced either as pre-cloned promoter-gene-terminator cassettes or as individual promoters, genes, and terminators containing 40 bp of overlap between parts. Integrations were targeted to a region of interest in *trans* through co-transformation of ∼600 bp regions of homology to the genome, with 40 bp of homology to common linker sequences present at the 5′ and 3′ ends of gene cassettes. All transformations were performed using a standard lithium acetate/salmon sperm heat shock protocol. Yeast strains were cured of Cas9-containing plasmids between rounds of transformations by sub-streaking on solid YPD.

The vector pGC1899, harboring expression cassettes for *SdiCFS*, *NdoSPS*, and *PsTNMT*, was constructed *via* Golden Gate assembly. Type IIS enzymes were purchased from Thermo Fisher Scientific, T7 ligase and T4 ligase buffer were purchased from New England Biolabs. Golden Gate reactions were performed as described in the Yeast Toolkit (YTK) system^19^. Assemblies containing *SdiCFS* were performed using “end on ligation”, due to the presence of an internal *Bsa*I site.

### Growth curves and determination of maximum growth rate

Yeast was grown in a Tecan Sunrise plate reader. Plates were wrapped with Parafilm to prevent evaporation. Growth measurements (OD_595_) were taken every 5 min for 48 hrs. Following background subtraction, values were normalized to starting OD, ln-transformed, and smoothed across a 20-minute interval. The slope of the curves across rolling 1-hour windows was determined, and the maximum slope was considered μ_max_.

### Fed-batch fermentation

Fed-batch fermentations were performed in Applikon 3L BioBundle fermenters. pH was maintained at 4.5 using 4N NaOH, temperature was kept at 30 °C. Air flow was set to 1 L/min, dissolved oxygen was controlled at 35% air saturation by automatic adjustment of stirring rate. Off-gas composition (partial pressure of O_2_ and CO_2_) was measured with a Tandem Multiplex gas analyzer. Precultures were grown at 30 °C for 36 hrs in 50 mL SC-His medium with shaking at 200 rpm. Cells were centrifuged for 10 min at 4000 *g*, washed in 0.9% NaCl, and suspended in 50 mL 0.9% NaCl prior to inoculation in 950 mL of batch medium (initial OD_600_∼0.2). Following exhaustion of sugar as indicated by off-gas analysis, fed-batch phase was triggered with an initial feeding rate of 0.60 g/h sucrose, increased exponentially at a dilution rate of 0.025 h^-1^. Batch medium (per liter): 40 g sucrose, 6 g (NH_4_)_2_SO_4_, 2.5 g/L KH_2_PO_4_, 1 g MgSO_4_·7H_2_O, 5 mL vitamin stock, and 5 mL trace element stock. Feeding medium (per liter): 360 g sucrose, 60 g (NH_4_)_2_SO_4_, 15 g KH_2_PO_4_, 6 g MgSO_4_·7H_2_O, 15 mL vitamin stock, and 15 mL trace element stock per liter. Vitamin stock (per liter): 2,500 mg myo-inositol, 100 mg calcium pantothenate, 100 mg thiamine hydrochloride, 100 mg pyridoxine, 100 mg nicotinic acid, 20 mg p-aminobenzoic acid, 5 mg biotin, and 5 mg folic acid. Trace element stock (per liter): 15 g Na_2_EDTA, 2.9 g CaCl_2_, 9.2 g ZnSO_4_·7H_2_O, 0.5 g CuSO_4_, 0.43 g MnSO_4_·H2O, 0.47 g CoCl_2_, 0.48 g Na_2_MoO_4_, and 5.1 g FeSO_4_·7H2O. A biomass conversion ratio of 0.59 g/L per OD_600_ unit was determined by drying and weighing cells in pre-dried Falcon tubes in a 100°C oven overnight in triplicate.

### High pressure liquid chromatography analysis by ultraviolet absorbance (HPLC-UV) and mass spectrometry (HPLC-MS)

Dopamine, hydroxytyrosol, tyrosol, 4-HPAC, 3,4-dHPAC, and (*S*)-reticuline were quantified by HPLC-UV using an Agilent 1200 HPLC system. Samples in 96-well plate format were diluted 1:2 with 100% acetonitrile (AcN) containing 0.1% trifluoroacetic acid (TFA), vortexed briefly, centrifuged for 5 min at 21,000 *g*, and then supernatant was analyzed. Supernatants from bioreactors were further diluted with 50% AcN/0.1% TFA as appropriate to stay within the range of standard curves. Five μL of analyte was applied to an Eclipse XDB-C18 column (150 x 4.6 mm, 5 μm, Agilent Technologies) and separated using the following gradient at 1 mL/min: 0-10 min, 5-20% B; 10-15 min, 20-50% B; 15-15.1 min, 50-95% B; 15.1-25 min, 95% B; 25-28 min, 5% B where A was 0.1% TFA in water and B was 0.1% TFA in methanol. All compounds were detected at 280 nm.

Relative peak areas of BIAs and BIA-like scaffolds were assessed using an Agilent 6545 qTOF-MS. All samples were diluted 1:5 with 100% AcN containing 0.1% formic acid (FA) and water containing 0.1% FA was added to bring the final AcN concentration to 15%. Samples were centrifuged for 10 min at 4,000 *g*, and then supernatant was analyzed. Supernatants were diluted as necessary to avoid saturation of the detector. Five μL of analyte was applied to a Zorbax Eclipse Plus C18 column (50 x 2.1 mm, 1.8 μm, Agilent Technologies) and separated using the following gradient at 0.3 mL/min flow rate: 0-4 min, 2-10% B; 4-6 min, 10-85% B; 6-7 min, 85% B, 7-7.1 min, 85-2% B where A was 0.1% FA in water and B was 0.1% FA in AcN. The column was reequilibrated for 2 min in 2% B at 0.45 ml/min. Settings: column compartment, 30°C; sheath gas flow rate, 10 L/min; sheath gas temperature, 350°C; drying gas flow rate 12 L/min; drying gas temperature, 325°C; nebulizing gas, 55 psig.

Dihydrosanguinarine pathway intermediates were assessed by HPLC-FT-MS using an Agilent 1290 Infinity II HPLC (Agilent Technologies) and a 7T-LTQ-FT-ICR (Thermo Fisher Scientific). Samples in 96-well plate format were extracted as per HPLC-qTOF-MS analysis. For determination of dihydrosanguinarine and sanguinarine concentration in bioreactors, fermentation broth was diluted 1:5 with 100% MeOH containing 0.1% HCl, and then further diluted to 1:100 with 100% MeOH/0.1% FA prior to centrifugation for 5 min at 4,000 *g*. Samples were diluted as necessary in 100% MeOH/0.1% FA to stay within the linear range of the MS. Five μL was applied to a Zorbax Eclipse Plus C18 column (50 x 2.1 mm, 1.8 μm, Agilent Technologies) and separated using the following gradient: 0.3 mL/min flow rate: 0-4 min, 2-10% B; 4-6 min, 10-85% B; 6-9 min, 85% B, 9-9.1 min, 85-2% B where A was 0.1% FA in water and B was 0.1% FA in AcN. The column was reequilibrated for 5 min in 2% B at 0.3 ml/min. Settings: scanning range, 100-400 *m/z*, resolution, 25,000; capillary voltage, 5 kV; source temperature, 350°C.

Sources of HPLC-UV/MS reagents: water and acetonitrile, Fisher Scientific; methanol, Sigma Aldrich; formic acid, Fluka; trifluoroacetic acid, Sigma Aldrich. Sources for authentic BIAs were: (*S*)-norcoclaurine, TRC Inc. (North York, Ontario, Canada); (*S*)-reticuline, gift from Dr. Peter Facchini; (*S*)-scoulerine, ChromaDex (Irvine, Ca, USA); (*S*)-stylopine, ChromaDex (Irvine, Ca, USA); protopine, TRC Inc.; sanguinarine, Sigma. Dihydrosanguinarine was derived from sanguinarine through sodium borohydride reduction^45^.

## Notes

### Competing Interest Statement

The authors have declared no competing interest.

