## Supplemental Information for "Benzylisoquinoline alkaloid production in yeast via norlaudanosoline improves selectivity and yield"

**Supplemental Figures**

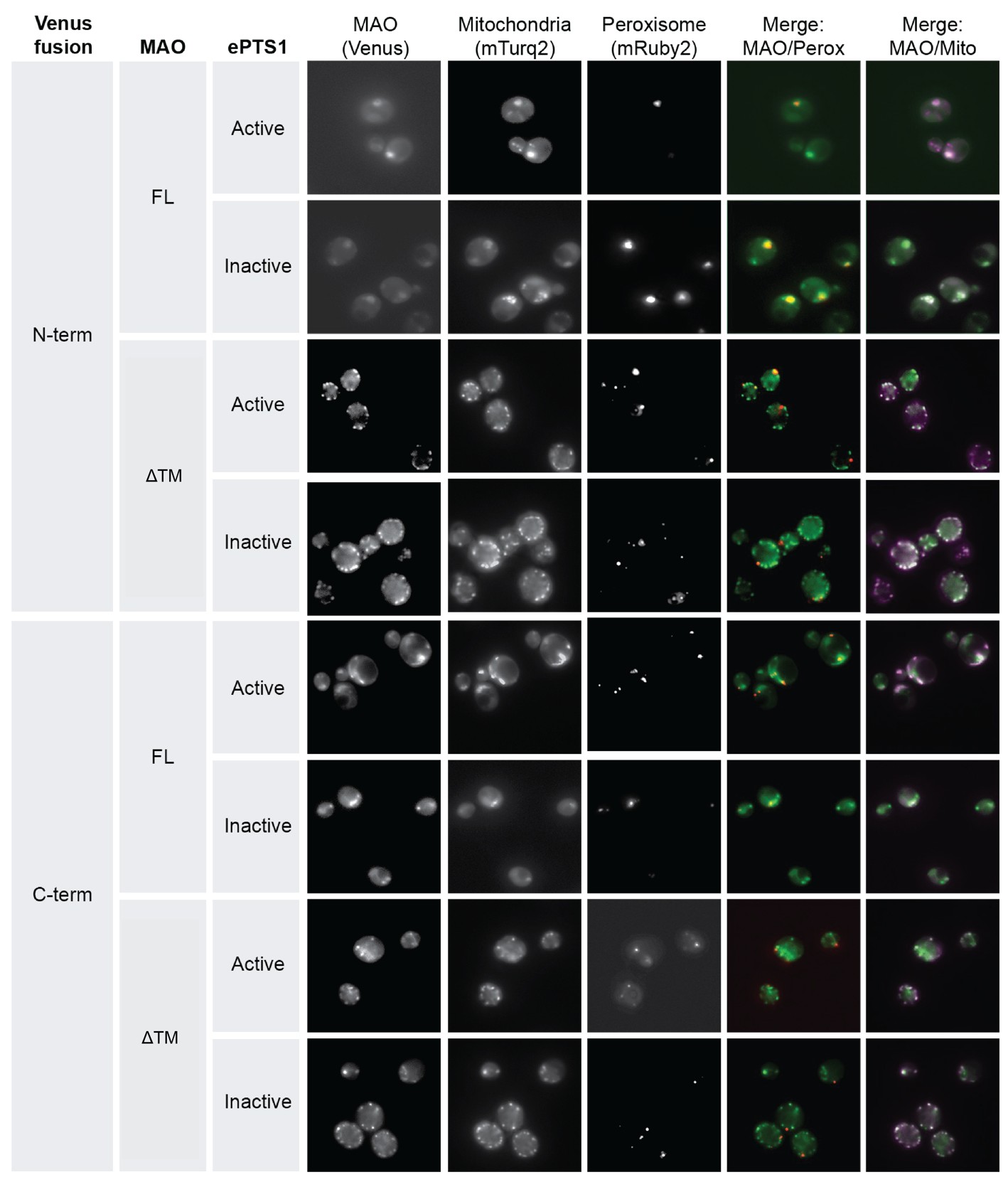

Supplemental Figure 1. Visualization of MAO localization by fluorescence microscopy

The fluorescent protein Venus was fused to the N-terminus (N-term) or C-terminus (C-term) of full-length MAO (FL) or MAOΔTM (ΔTM). At the extreme C-terminus of each fusion protein was an active ePTS1, targeting the cargo to the peroxisome, or an inactive ePTS1, expected to maintain cytosolic localization. Each fusion protein was expressed in strain yKSS001, which harbors *Su9ss-mTurq2* to visualize mitochondria and *mRuby-ePTS1* to visualize peroxisomes. Although mitochondrial morphology differs between strains expressing *MAO* and *MAOΔTM,* mitochondrial co-localization is observed throughout*.* Further, no differences are observed between MAO variants with active or inactive ePTS1.

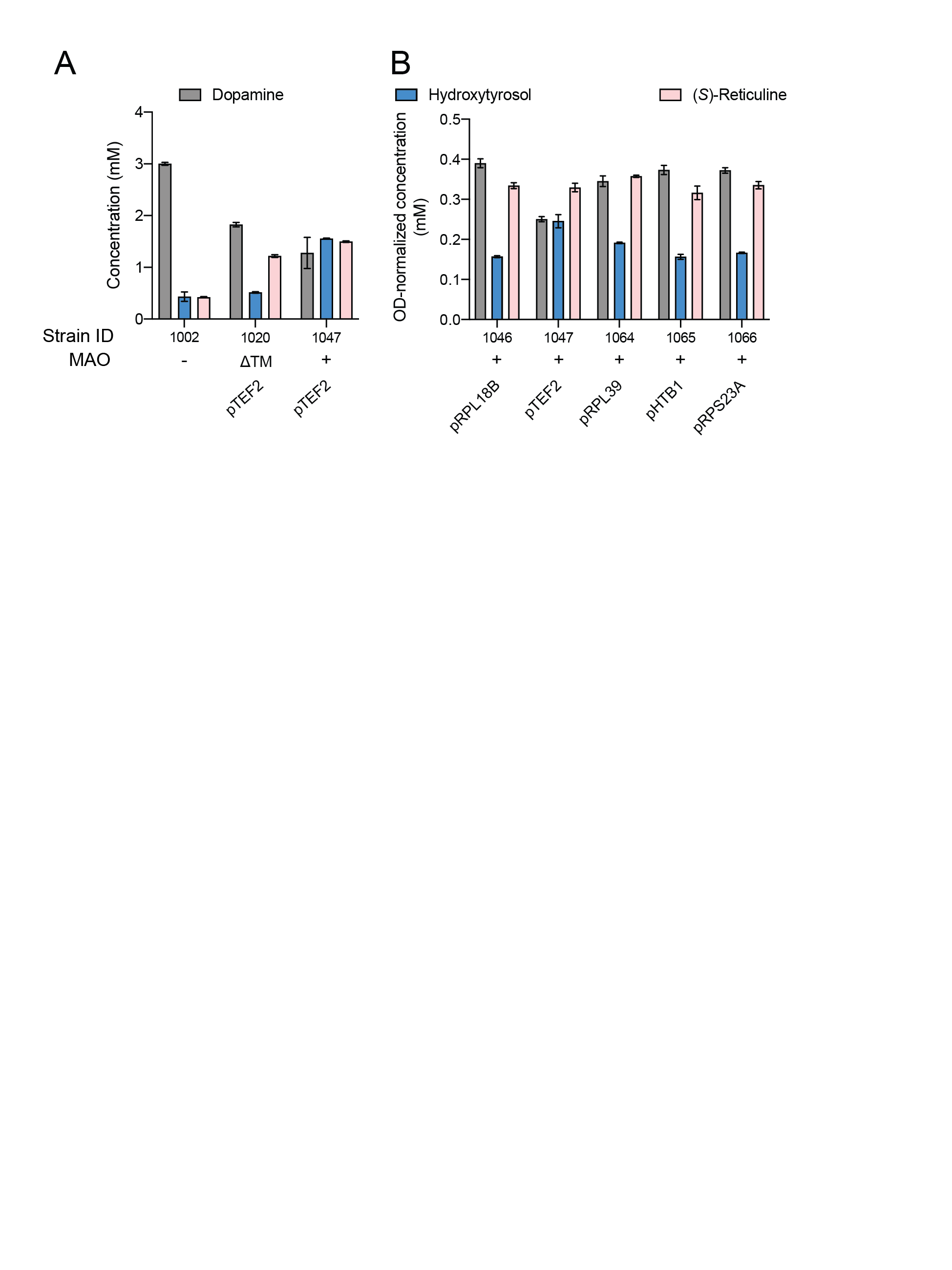

Supplemental Figure 2. Metabolite profile of strains expressing *MAO*

*MAO*, with (+) or without its C-terminal transmembrane helix (ΔTM), was integrated into strain LN1002 (*aro10*Δ) under a variety of promoter strengths. Strains were grown in synthetic complete media in deep well plates and then analyzed for dopamine, reticuline, and hydroxytyrosol content by LC-UV. Error bars refer to the mean and standard deviation of n=2 biological replicates. **(A)** Strain LN1020, harboring MAOΔTM, produces more dopamine, less hydroxytyrosol, and less reticuline than strain LN1047, harboring full-length MAO. **(B)** Strains LN1064, LN1065, and LN1066, expressing *MAO* from the promoters of *RPL39*, *HTB1*, and *RPS23A*, respectively, have a metabolite profile that resembles strains LN1046 and LN1047. Metabolite content is represented normalized to final OD_600_.

­­
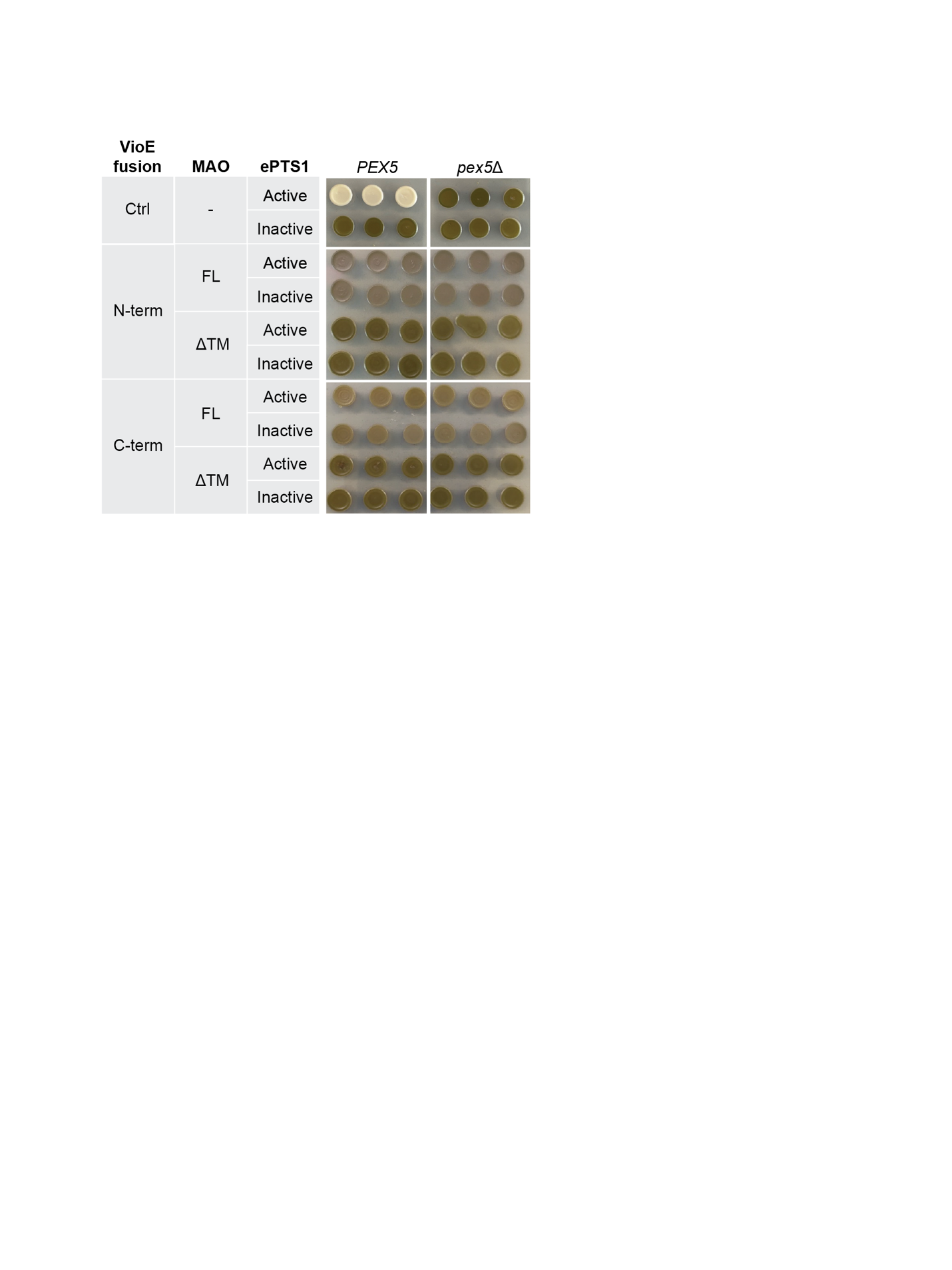

Supplemental Figure 3. Assessment of MAO peroxisomal compartmentalization

Agar spot assay to visualize efficiency of MAO import to the peroxisome. Strains yPSG335 (*PEX5*) and yPSG340 (*pex5*Δ) express the violacein pathway proteins VioA and VioB in the cytosol, which convert tryptophan to IPA imine dimer (colorless). VioE, fused to MAO or MAOΔTM at the N- or C-terminus, is required to convert IPA imine dimer to prodeoxyviolacein (green). Sequestration of VioE to the peroxisome, *via* active ePTS1, prevents the formation of colour (Ctrl, active ePTS1, *PEX5*). Inactive ePTS1 reduces peroxisomal import dramatically^1^; such strains should be a deep green (Ctrl, inactive ePTS1, *PEX5*). *pex5*Δ strains, defective in peroxisomal import, are also deep green. A lack of difference in colour between active and inactive ePTS1 fusion proteins indicates that MAO peroxisomal import is not effective. Expression of full-length MAO, especially as a VioE N-terminal fusion, results in a grey colour. Abbreviations: ePTS1, enhanced peroxisomal targeting sequence 1; Ctrl, control; FL, full-length; ΔTM, MAOΔTM; N-term, N-terminal enzyme fusion; C-term, C-terminal enzyme fusion.

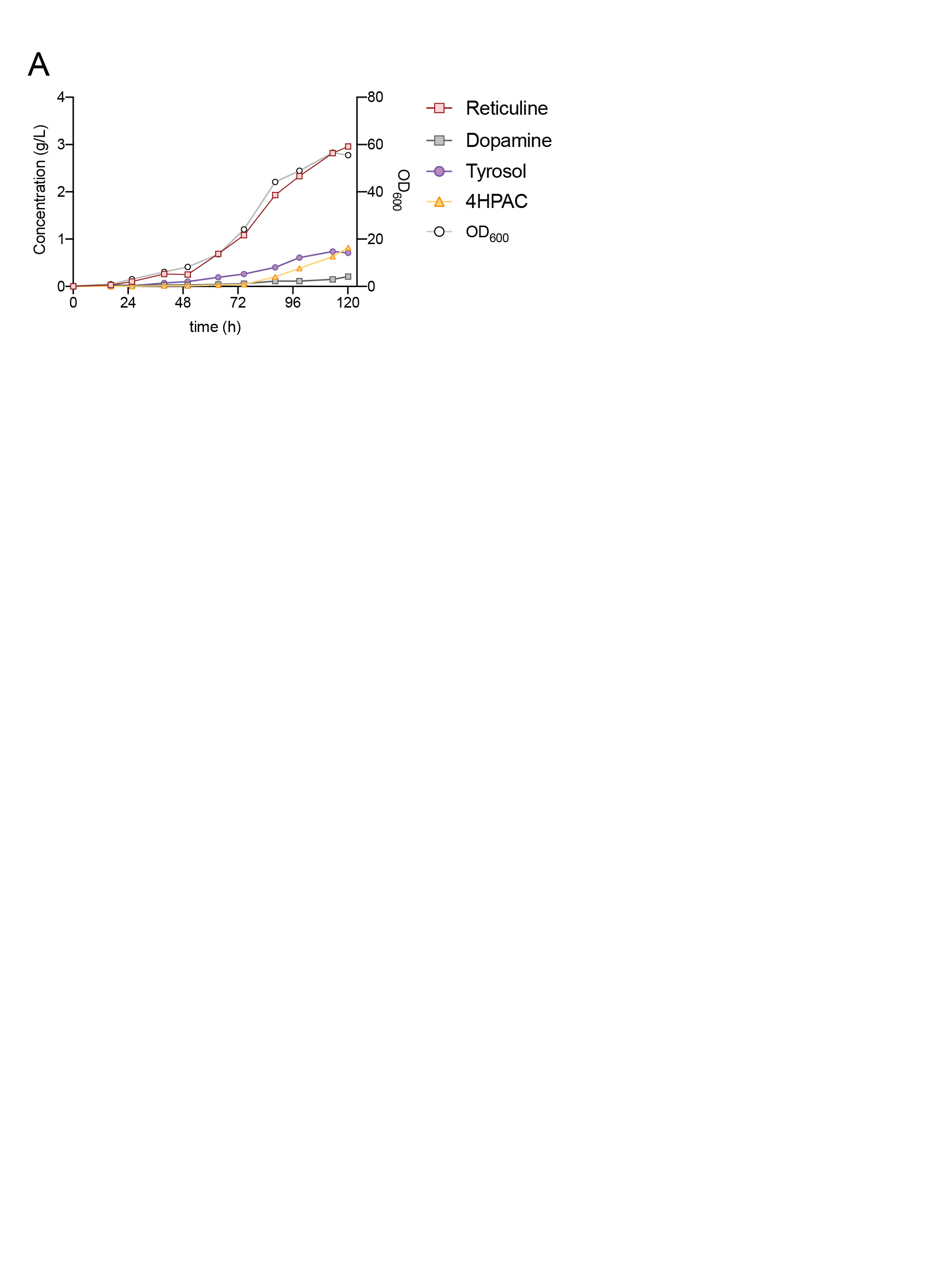

Supplemental Figure 4. Fed-batch fermentation of strain LP524

Strain LP524 was grown in a bioreactor using an exponential, sugar-limited feeding profile. Samples were regularly taken and assessed for biomass and for metabolite profile by LC-UV.

**Supplemental Materials & Methods**

*Strain construction and growth media for MAO localization experiments*

The *S. cerevisiae* strains for MAO localization and visualization experiments were BY4741 (*MATa his3Δ1 leu2Δ0 met15Δ0 ura3Δ0*) and BY4741 *pex5Δ*, ordered from Open Biosystems (GE Dharmacon). Wild-type yeast cultures were grown in YPD medium (10 g/L Bacto yeast extract, 20 g/L Bacto peptone, 20 g/L glucose). Selection of auxotrophic markers (*URA3*, *LEU2*, *HIS3*) was performed in synthetic complete (SC) medium (6.7 g/L Difco Yeast Nitrogen Base without amino acids (Spectrum Chemical); 2 g/L Drop-out Mix Synthetic minus appropriate amino acids, without Yeast Nitrogen Base (US Biological); 20 g/L glucose). All strains used are listed in Table 4.1

Golden Gate assembly reactions were transformed into chemically competent *E. coli* prepared from strain TG1 (Lucigen). Transformed cells were selected on LB containing the antibiotics chloramphenicol, ampicillin or kanamycin.

Yeast expression vectors were built using Golden Gate assembly as described in the Yeast Toolkit (YTK) system^2^. Integration into the yeast genome via homologous recombination at the *URA3* or *LEU2* locus was achieved by transformation of linearized plasmids (*Not*I digestion, NEB). All transformations were performed using a standard lithium acetate method, and cells were plated onto selective auxotrophic SC agar plates with 2% glucose. Individual colonies were picked as biological replicates directly from this transformation plate and grown independently for further analysis.

Chromosomal integrations of the fluorescent localization reporters mRuby2-ePTS1 (peroxisome) and Su9(1-69)-mTurqoise2^3^ (mitochondria) were performed by co-transforming a CEN6/ARS4 CRISPR plasmid (containing Cas9, a guide RNA for targeting the YMR206W locus and a *HIS3* auxotrophic marker) with linearized repair DNA designed for homologous recombination. Cells were plated on histidine-dropout medium, re-streaked on histidine-dropout medium and then grown in non-selective medium to remove the CRISPR plasmid. Chromosomal integration was confirmed by PCR, and removal of CRISPR plasmid was confirmed by replica plating from non-selective medium onto histidine-dropout medium (a colony will not grow on histidine-dropout medium if the CRISPR plasmid has been removed).

*Fluorescence microscopy*

For MAO localization experiments, strains were grown to saturation in SC medium with 2% glucose and auxotrophic selection, diluted 50-fold into fresh selective medium and grown for 6–8 hrs. Cultures were resuspended in 1x PBS before spotting onto plain glass slides for imaging on a Zeiss Axio Observer D1 microscope with an X-Cite Series 120 fluorescent lamp, a Hamamatsu Orca-Flash 4.0 digital camera and ZEN 2.6 (blue edition) software. Images were analyzed using ZEN 3.0 (blue edition) software. The fluorescent protein variants used were CFP mTurqoise2, YFP Venus and RFP mRuby2.

*Spot assays for prodeoxyviolacein production and visualization*

Agar plate spots for visualization of prodeoxyviolacein production were generated by plating 5 µl of saturated culture on SC agar plates with 2% glucose and appropriate auxotrophic selection. Plates were grown at 30 °C and imaged at 48 hrs. Three biological replicates of each strain were plated and imaged.

**Supplemental tables**

Table S1. Strain lists

| **Base strain for BIA synthesis** | | | |
| --- | --- | --- | --- |
| **Name** | **Brief description** | **Genotype** | **Ref.** |
| LP507 | Strain derived from BY4741 capable of making 4.6 g/L reticuline. See reference for full detail. | *BY4741*. Oxidoreductase knockouts: Δ*aad3*, Δ*ydr541c*, Δ*ypr1*, Δ*adh6*, Δ*ari1*, Δ*gre2*, Δ*hfd1*. Mitochondrial fixes: *MIP1*, *CAT5*, *SAL1*. Aromatic amino acid flux: *ARO4^FBR^*, *ARO7^FBR^*, *ARO2*, *TYR1*. 4-HPAA synthesis: *ARO10*. Dopamine synthesis: *DODC*, *CYP76AD5*. Norcoclaurine synthesis: *CjNCSΔ35* (2 copies). Reticuline pathway: *Ps6OMT*, *PsCNMT*, *Ps4′OMT* (2 copies), *AtATR2*. | ^4^ |

| **Yeast strains generated for BIA synthesis** | | | |
| --- | --- | --- | --- |
| **Name** | **Parent** | **Brief Description** | **Genotype** |
| LP524 | LP507 | Complete second copy of reticuline pathway | **USERXII-1**::P_TDH3_-*Ps*CNMT-T_TDH1_-  P_CCW12_-*Ec*NMCH-T_PGI1_ |
| LN1002 | LP524 | Knockouts of native and overexpressed Aro10 | Δ*Aro10*, Δ**FgF18**::P_TDH3_-Aro10-T_TDH1_ |
| LN1044 | LN1002 | RNR2-MAO | **USERX-1**::P_RNR2_-*Hs*MAO-A-T_TDH1_ |
| LN1046 | LN1002 | RPL18B-MAO | **USERX-1**::P_RPL18B_-*Hs*MAO-A-T_TDH1_ |
| LN1047 | LN1002 | TEF2-MAO | **USERX-1**::P_TEF2_-*Hs*MAO-A-T_TDH1_ |
| LN1049 | LN1002 | TDH3-MAO | **USERX-1**::P_TDH3_-*Hs*MAO-A-T_TDH1_ |
| LN1020 | LN1002 | TEF2-MAOΔTM | **USERX-1**::P_TEF2_-*Hs*MAO-AΔTM-T_TDH1_ |
| LN1015 | LN1002 | TEF2-MAOΔTM-p | **USERX-1**::P_TEF2_-*Hs*MAO-AΔTM-ePTS1-T_TDH1_ |
| LN1063 | LN1046 | NCS-p | **USERX-3**::P_TDH3_-*Cj*NCSΔ35-ePTS1-T_ENO2_ |
| LN1064 | LN1002 | RPL39-MAO | **USERX-1**::P_RPL39_-*Hs*MAO-A-T_TDH1_ |
| LN1065 | LN1002 | HTP1-MAO | **USERX-1**::P_HTB1_-*Hs*MAO-A-T_TDH1_ |
| LN1066 | LN1002 | RPS23A-MAO | **USERX-1**::P_RPS23A_-*Hs*MAO-A-T_TDH1_ |
| LN1067 | LN1064 | NCS-p | **USERXI-2**::P_TDH3_-*Cj*NCSΔ35-ePTS1-T_ENO2_ |
| LN1068 | LN1065 | NCS-p | **USERXI-2**::P_TDH3_-*Cj*NCSΔ35-ePTS1-T_ENO2_ |
| LN1069 | LN1066 | NCS-p | **USERXI-2**::P_TDH3_-*Cj*NCSΔ35-ePTS1-T_ENO2_ |
| LN1075 | LN1015 | BBE | **USERX-3**::P_TDH3_-*Ec*BBE-T_TDH1_ |
| LN1078 | LN1075 | CFS/SPS/TNMT | **USERXI-2**::P_PGK1_-*Sdi*CFS-T_ADH1_-P_TEF1_-*Ndo*SPS-T_PGK1_-P_TEF2_-TNMT-T_ADH1_ |
| LN1079 | LN1075 | BBE (2 copies) | **USERXI-2**::P_TDH3_-*Ec*BBE-T_TDH1_ |
| LN1080 | LN1079 | CFS/SPS/TNMT | **USERIV-1**:: P_PGK1_-*Sdi*CFS-T_ADH1_-P_TEF1_-*Ndo*SPS-T_PGK1_-P_TEF2_-TNMT-T_ADH1_ |
| LN1083 | LN1080 | MSH/P6H | **USERIX-1**::PPDC1-P6H-TCYC1-**linker**-PTDH3-MSH-TADH1 |
| LN1084 | LN1080 | MSH/P6H/TNMT | **USERIX-1**::PPDC1-P6H-TCYC1-**linker**-PTDH3-MSH-TADH1-**linker**-PFBA1-TNMT-TPGI1 |

| **Yeast strains generated for assessment of MAO localization** | | | |
| --- | --- | --- | --- |
| **Name** | **Parent** | **Plasmid used** | **Description** |
| yWCD230 | BY4741 | pWCD1351 | Wild-type yeast (*his3*Δ) |
| yWCD231 | BY4741 pex5Δ::KanMX | pWCD1351 | Defective peroxisomal import (*his3*Δ) |
| yKSS001 | yWCD230 | pKS0864 | YMR206WΔ::pHHF1-Su9(1-69)-mTurquoise2-tENO2-pTEF2-mRuby2-GSLGRGRR-SKL!-tPGK1 |
| yKSS002 | yKSS001 | pKS0902 | Microscopy: N-terminal VioE+Venus MAO, ePTS1 |
| yKSS003 | yKSS001 | pKS0903 | Microscopy: N-terminal VioE+Venus MAO, inactive ePTS1 |
| yKSS004 | yKSS001 | pKS0904 | Microscopy: N-terminal VioE+Venus MAOΔTM, ePTS1 |
| yKSS005 | yKSS001 | pKS0905 | Microscopy: N-terminal VioE+Venus MAOΔTM, inactive ePTS1 |
| yKSS006 | yKSS001 | pKS0906 | Microscopy: C-terminal VioE+Venus MAO, ePTS1 |
| yKSS007 | yKSS001 | pKS0907 | Microscopy: C-terminal VioE+Venus MAO, inactive ePTS1 |
| yKSS008 | yKSS001 | pKS0908 | Microscopy: C-terminal VioE+Venus MAOΔTM, ePTS1 |
| yKSS009 | yKSS001 | pKS0909 | Microscopy: C-terminal VioE+Venus MAOΔTM, inactive ePTS1 |
| yPSG335 | yWCD230 | pJAS1052 | pTDH3-VioA-tENO1-pTEF1-VioB-tPGK1 |
| yPSG340 | yWCD231 | pJAS1052 | pTDH3-VioA-tENO1-pTEF1-VioB-tPGK1 |
| yKSS010 | yPSG335 | pKS0902 | Violacein: N-terminal VioE+Venus MAO, ePTS1 |
| yKSS011 | yPSG335 | pKS0903 | Violacein: N-terminal VioE+Venus MAO, inactive ePTS1 |
| yKSS012 | yPSG335 | pKS0904 | Violacein: N-terminal VioE+Venus MAOΔTM, ePTS1 |
| yKSS013 | yPSG335 | pKS0905 | N-terminal VioE+Venus MAOΔSP, inactive ePTS1 |
| yKSS014 | yPSG335 | pKS0906 | Violacein: C-terminal VioE+Venus MAO, ePTS1 |
| yKSS015 | yPSG335 | pKS0907 | Violacein: C-terminal VioE+Venus MAO, inactive ePTS1 |
| yKSS016 | yPSG335 | pKS0908 | Violacein: C-terminal VioE+Venus MAOΔTM, ePTS1 |
| yKSS017 | yPSG335 | pKS0909 | Violacein: C-terminal VioE+Venus MAOΔTM, inactive ePTS1 |
| yKSS018 | yPSG340 | pKS0902 | Violacein, import defects: N-terminal VioE+Venus MAO, ePTS1 |
| yKSS019 | yPSG340 | pKS0903 | Violacein, import defects: N-terminal VioE+Venus MAO, inactive ePTS1 |
| yKSS020 | yPSG340 | pKS0904 | Violacein, import defects: N-terminal VioE+Venus MAOΔTM, ePTS1 |
| yKSS021 | yPSG340 | pKS0905 | Violacein, import defects: N-terminal VioE+Venus MAOΔTM, inactive ePTS1 |
| yKSS022 | yPSG340 | pKS0906 | Violacein, import defects: C-terminal VioE+Venus MAO, ePTS1 |
| yKSS023 | yPSG340 | pKS0907 | Violacein, import defects: C-terminal VioE+Venus MAO, inactive ePTS1 |
| yKSS024 | yPSG340 | pKS0908 | Violacein, import defects: C-terminal VioE+Venus MAOΔTM, ePTS1 |
| yKSS025 | yPSG340 | pKS0909 | Violacein, import defects: C-terminal VioE+Venus MAOΔTM, inactive ePTS1 |
| yKSS026 | yPSG335 | pKS1124 | Violacein: VioE-Venus fusion, ePTS1 |
| yKSS027 | yPSG335 | pKS1125 | Violacein: VioE-Venus fusion, inactive ePTS1 |
| yKSS028 | yPSG340 | pKS1124 | Violacein, import defects: VioE-Venus fusion, active ePTS1 |
| yKSS029 | yPSG340 | pKS1125 | Violacein, import defects: VioE-Venus fusion, inactive ePTS1 |

Table S2. Plasmid list

| **Plasmids** | | |
| --- | --- | --- |
| **Name** | **Brief Description** | **Ref.** |
| pCAS-G418 | Yeast replicative plasmid (2μ) with Cas9 and tRNA^Tyr^-driven gRNA. G418 resistance. | ^5^ |
| pCAS-Hyg | Yeast replicative plasmid (2μ) with Cas9 and tRNA^Tyr^-driven gRNA. Hyg resistance. | ^4^ |
| pBSC011 | P_TDH3_-CjNCSΔ35-T_ENO2_ | ^4^ |
| pGC1899 | Harbors CFS, SPS, TNMT for integration into genome | This work |
| pGC997 | Harbors MSH, P6H, TNMT for integration into genome | ^6^ |
| pHUM | Harbors *HIS3*, *URA3*, *MET17* for prototrophy restoration of S288C-derived strains | ^7^ |
| pWCD1351 | *HIS3* | This work |
| pKS0864 | Integrative plasmid harbouring pHHF1-Su9ss-mTurquoise2-tENO2-pTEF2-mRuby2-GSLGRGRR-SKL!-tPGK1 and a *HIS3* marker | This work |
| pKS0902 | pTEF2-VioE-Venus-Hs_MAO-A-ePTS1-tADH1 | This work |
| pKS0903 | pTEF2-VioE-Venus-Hs_MAO-A-dead_PTS1-tADH1 | This work |
| pKS0904 | pTEF2-VioE-Venus-Hs_MAO-A(1-497)-ePTS1-tADH1 | This work |
| pKS0905 | pTEF2-VioE-Venus-Hs_MAO-A(1-497)-dead_PTS1-tADH1 | This work |
| pKS0906 | pTEF2-Hs_MAO-A-VioE-Venus-ePTS1-tADH1 | This work |
| pKS0907 | pTEF2-Hs_MAO-A-VioE-Venus-dead_PTS1-tADH1 | This work |
| pKS0908 | pTEF2-Hs_MAO-A(1-497)-VioE-Venus-ePTS1-tADH1 | This work |
| pKS0909 | pTEF2-Hs_MAO-A(1-497)-VioE-Venus-dead_PTS1-tADH1 | This work |
| pKS1124 | pTEF2-VioE-Venus-ePTS1-tADH1 | This work |
| pKS1125 | pTEF2-VioE-Venus-dead_PTS1-tADH1 | This work |

Table S3. Primer list

| **Yeast strains** | | | |
| --- | --- | --- | --- |
| **Number** | **Name** | **Sequence (5′->3′)** | **Description** |
| LB33 | HDVgRNA_F | CACCTATATCTGCGTGTTGC | When paired: amplify empty guide RNA transcription cassette from pCAS-G418/Hyg. Use with two internal primers providing guide RNA target sequence. |
| LB34 | gRNA_scaffold_R | GTCAAGACTGTCAAGGAGG |  |
| LB1968 | X-1_UP_F | cgctcactagtagacaacacacg | Amplify X-1 upstream homology region with homology to gene cassettes |
| LB1969 | (LV3)_X-1_UP_R | gcatttttattatataagttgttttattcagagtattcctggaagcatacagaatattcactaac |  |
| LB1970 | (LV5)_X-1_DN_F | Cctctttatattacatcaaaataagaaaataattataacaccacgattgagtgtctgcactcttattc | Amplify X-1 downstream homology region with homology to gene cassettes |
| LB1971 | X-1_DN_R | cctttttccaattcttaggctatttgg |  |
| LB1964 | X-1_guide_F | gtagctacaagaacatatggGTTTTAGAGCTAGAAATAGCAAGT | Use with LB33/LB34 to introduce X-1 guide sequence to CRISPR plasmid |
| LB1965 | X-1_guide_R | ccatatgttcttgtagctacAAAGTCCCATTCGCCACCCGAA |  |
| LB1987 | X-3_UP_F | ggctactgattttgttaagcaactc | Amplify X-3 upstream homology region with homology to gene cassettes |
| LB1979 | (LV3)_X-3_UP_R | gcatttttattatataagttgttttattcagagtattcctgagaagaaattttgggggtaatatg |  |
| LB1980 | (LV5)_X-3_DN_F | Cctctttatattacatcaaaataagaaaataattataacagggaaataaggtttaaaggcactg | Amplify X-3 downstream homology region with homology to gene cassettes |
| LB1981 | X-3_DN_R | ggtatctcaatgaacgagctcg |  |
| LB1974 | X-3_guide_F | gatcgccgaatggcacgcgaGTTTTAGAGCTAGAAATAGCAAGT | Use with LB33/LB34 to introduce X-3 guide sequence to CRISPR plasmid |
| LB1975 | X-3_guide_R | tcgcgtgccattcggcgatcAAAGTCCCATTCGCCACCCGAA |  |
| LB1988 | XI-2_UP_F | ggtttctgaaaaaagaagtagtcg | Amplify XI-2 upstream homology region with homology to gene cassettes |
| LB1989 | (LV3)_XI-2_UP_R | gcatttttattatataagttgttttattcagagtattcctctgaaagcgctagtcgtgtgtacc |  |
| LB1990 | (LV5)_XI-2_DN_F | Cctctttatattacatcaaaataagaaaataattataacagctttgcagttttcgtggctag | Amplify XI-2 downstream homology region with homology to gene cassettes |
| LB1991 | XI-2_DN_R | ctataacatggtttacaaacccgagg |  |
| LB1984 | XI-2_guide_F | actttgtcgtttcttactttGTTTTAGAGCTAGAAATAGCAAGT | Use with LB33/LB34 to introduce XI-2 guide sequence to CRISPR plasmid |
| LB1985 | XI-2_guide_R | aaagtaagaaacgacaaagtAAAGTCCCATTCGCCACCCGAA |  |
| LN1100 | IV-1_UP_F | CGTGCGCTTGAGATTCAGT | Amplify IV-1 upstream homology region with homology to gene cassettes |
| LN1101 | (LV3)_IV-1_UP_R | gcatttttattatataagttgttttattcagagtattcctAGAGTTCCCGTCGGAAT |  |
| LN1102 | (LV5)_IV-1_DN_F | cctctttatattacatcaaaataagaaaataattataacaCGTTACTAGCGTTGCAAGTGG | Amplify IV-1 downstream homology region with homology to gene cassettes |
| LN1103 | IV-1_DN_R | GGATTTGGTTTAGCAGCAGTC |  |
| LN1106 | IV-1_guide_F | CGCCGGCTGGGCAACACCTTCGGGTGGCGAATGGGACTTTCTGCAAGGAAGTTTAAGCGT | Introduce IV-1 guide RNA sequence to CRISPR plasmid using overlap extension |
| LN1107 | IV-1_guide_R | GACTAGCCTTATTTTAACTTGCTATTTCTAGCTCTAAAACACGCTTAAACTTCCTTGCAGA |  |
| LN1092 | II-1_UP_F | GCGTTCACAGTTACTCTTTTAGAAC | Amplify II-1 upstream homology region with homology to gene cassettes |
| LN1093 | (LV3)_II-1_UP_R | gcatttttattatataagttgttttattcagagtattcctAAAATAACATGTTGCGTGCAC |  |
| LN1094 | (LV5)_II-1_DN_F | cctctttatattacatcaaaataagaaaataattataacaGACAAACTTTACAAAGAAGACACCC | Amplify II-1 downstream homology region with homology to gene cassettes |
| LN1095 | II-1_DN_R | GTATGCCGTGATATGAACAAAC |  |
| LN1098 | II-1_guide_F | CGCCGGCTGGGCAACACCTTCGGGTGGCGAATGGGACTTTAACTGCTCAGGGCGGATAAC | Introduce II-1 guide RNA sequence to CRISPR plasmid using overlap extension |
| LN1099 | II-1_guide_R | GACTAGCCTTATTTTAACTTGCTATTTCTAGCTCTAAAACGTTATCCGCCCTGAGCAGTT |  |
| LN1108 | IX-1_UP_F | CAACTGCTAAGAACTCTGTGATCTTC | Amplify IX-1 upstream homology region with homology to gene cassettes |
| LN1109 | (LV3)_IX-1_UP_R | gcatttttattatataagttgttttattcagagtattcctTCGCGAGATAGAACGACATC |  |
| LN1110 | (LV5)_IX-1_DN_F | cctctttatattacatcaaaataagaaaataattataacaTTGATGACACTAGCGGACTTG | Amplify IX-1 downstream homology region with homology to gene cassettes |
| LN1111 | IX-1_DN_R | GGCAGAAAACTACCCGTAGAATAC |  |
| LN1114 | IX-1_guide_F | CGCCGGCTGGGCAACACCTTCGGGTGGCGAATGGGACTTTATCTTAAATGAAAGACAGAGGTTTT | Introduce IX-1 guide RNA sequence to CRISPR plasmid using overlap extension |
| LN1115 | IX-1_guide_R | GACTAGCCTTATTTTAACTTGCTATTTCTAGCTCTAAAACCTCTGTCTTTCATTTAAGATAAAG |  |
| LN14 | (LV3)_pRNR2_F | aggaatactctgaataaaacaacttatataataaaaatgcAGTCGAACAAGAAGCAGG | Amplify pRNR2 with homology to gene cassettes |
| LN456 | pRNR2_R | GGTAATTGGACAAATAAATACGTGTATTAAG |  |
| LN18 | (LV3)_pRPL18B_F | aggaatactctgaataaaacaacttatataataaaaatgcaagaggatgtccaatattttt | Amplify pRPL18B with homology to gene cassettes |
| LN455 | pRPL18B_R | tttgttttttgttttcttctaattgatt |  |
| LN1080 | (LV3)_pRPL39_F | aggaatactctgaataaaacaacttatataataaaaatgcCTTGGATATGTATGTTGGTCTTGTT | Amplify pRPL39A with homology to gene cassettes |
| LN1081 | pRPL39_R | GTTGATCTATCTGTGTTTATTTGCTTG |  |
| LN1083 | (LV3)_pHTB1_F | aggaatactctgaataaaacaacttatataataaaaatgcATGATGGTTCAACAAGACCAGA | Amplify pHTB1 with homology to gene cassettes |
| LN1084 | pHTB1_R | TGTATGTGTGTATGGTTTATTTGTGG |  |
| LN1086 | (LV3)_pRPS23A_F | aggaatactctgaataaaacaacttatataataaaaatgcGTCGGTCGCACTAGACTTTTC | Amplify pRPS23A with homology to gene cassettes |
| LN1087 | pRPS23A_R | CTTTGTTTATTTCCTGTTGTCTTAGG |  |
| LN27 | (LV3)_pTEF2_F | aggaatactctgaataaaacaacttatataataaaaatgcttgataggtcaagatcaatgtaaac | Amplify pTEF2 with homology to gene cassettes |
| LN454 | pTEF2_R | gtttagttaattatagttcgttgaccg |  |
| PP120 | (LV3)_pTDH3_F | AGGAATACTCTGAATAAAACAACTTATATAATAAAAATGCtcgagtttatcattatcaatact | Amplify pTDH3 with homology to gene cassettes |
| LN398 | pTDH3_R | TTTGTTTGTTTATGTGTGTTTATtcg |  |
| LB1130 | tTDH1_F | ATAAAGCAATCTTGATGAGGATAATG | Amplify tTDH1 with homology to gene cassettes |
| LB1131 | (LV5)_tTDH1_R | tgttataattattttcttattttgatgtaatataaagagGCCATCCTAGAACTTCAATTCACCAC |  |
| LN247 | tPGI1_F | aacaaatcgctcttaaatatatacc | Amplify tPGI1 with homology to gene cassettes |
| PP127 | (LV5)_tPGI1_R | tgttataattattttcttattttgatgtaatataaagaggggtatactggaggcttcat |  |
| LN1018 | Aro10g1_OE_F | CGCCGGCTGGGCAACACCTTCGGGTGGCGAATGGGACTTTATATTCGCCTTGTGGACACT | Introduce guide RNA targeting Aro10 to CRISPR plasmid using overlap extension |
| LN1019 | Aro10g1_OE_R | GACTAGCCTTATTTTAACTTGCTATTTCTAGCTCTAAAACAGTGTCCACAAGGCGAATAT |  |
| LN1020 | Aro10_repair_OE_F | ATTGCCGAGGTCATGCTGAGCATTTGTCGTACTTTTGTGCCGTATATTAAAG | Generate repair template for native Aro10 knockout using overlap extension |
| LN1021 | Aro10_repair_OE_R | AAAGAACTCTGTGGTAGTGGTAAAATAGCTTTAATATACGGCACAAAAGTACGA |  |
| LN1022 | FgF18_repair_OE_F | tttcagtagatttggtaactgtgcaaccataactcatgccaatcgtc | Generate repair template for heterologous Aro10 knockout using overlap extension |
| LN1023 | FgF18_repair_OE_R | tgtgatgaattttgagagcccacttttgttggggacgattggcatgagttatg |  |
| LN459 | (pRNR2)_HsMAO-A_F | GAATCCAAACTTAATACACGTATTTATTTGTCCAATTACCATGGAAAATCAAGAAAAGGCATC | Amplify *Hs*MAO-A with homology to pRNR2 |
| LN458 | (pRPL18B)_HsMAO-A_F | atagaaagaaaaaatcaattagaagaaaacaaaaaacaaaATGGAAAATCAAGAAAAGGCATC | Amplify *Hs*MAO-A with homology to pRPL18B |
| LN1089 | (pRPL39)_HsMAO-A-F | AATTCGAAAAAGACAAGCAAATAAACACAGATAGATCAACATGGAAAATCAAGAAAAGGCATC | Amplify *Hs*MAO-A with homology to pRPL39 |
| LN1090 | (pHTB1)_HsMAO-A_F | ATAGACAAGTCAAACCACAAATAAACCATACACACATACAATGGAAAATCAAGAAAAGGCATC | Amplify *Hs*MAO-A with homology to pHTB1 |
| LN1091 | (pRPS23A)_HsMAO-A_F | AAATTTTACAAAAACCTAAGACAACAGGAAATAAACAAAGATGGAAAATCAAGAAAAGGCATC | Amplify *Hs*MAO-A with homology to pRPS23A |
| LN457 | (pTEF2)_HsMAO-A_F | tttttagaatatacggtcaacgaactataattaactaaacATGGAAAATCAAGAAAAGGCATC | Amplify *Hs*MAO-A with homology to pTEF2 |
| LN396 | (pTDH3)_HsMAO-A_F | caagaacttagtttcgaATAAACACACATAAACAAACAAAATGGAAAATCAAGAAAAGGCATC | Amplify *Hs*MAO-A with homology to pTDH3 |
| LN397 | (tTDH1)_HsMAO-A_FL_R | attcaaaaaaaaatcattatcctcatcaagattgctttatTTATGATCTTGGCAACAATTTGTAC | Amplify full-length *Hs*MAO-A with homology to tTDH1 |
| LN978 | (ePTS1)_HsMAO-A_dSP_R | CAACTTAGAACGACGACCACGACCTAATGATGGCAAGTTTCTTTCCCAAAAaGTATGAGTGATTTCAAC | Amplify signal peptide-less *Hs*MAO-A with C-terminal ePTS1 tag |
| LN979 | (ePTS1)_TDH1t_F | TTAGGTCGTGGTCGTCGTTCTAAGTTGTAAataaagcaatcttgatgaggataatg | Amplify tTDH1 with homology to C-terminal ePTS1 tag |
| LN1071 | (pTEF1)_CjNCS_F | cttcttgctcattagaaagaaagcatagcaatctaatctaagttttaatAAACAAtggaagaaactgtaatgttatatc | Amplify *Cj*NCS with homology to pTEF1 |
| LN1072 | (ePTS1)_CjNCS_R | ttaCAACTTAGAACGACGACCACGACCTAActcagaagatttgtgcttattttc | Amplify *Cj*NCS with C-terminal ePTS1 tag |
| LN1070 | (ePTS1)_tPGI1_F | TTAGGTCGTGGTCGTCGTTCTAAGTTGtaaaacaaatcgctcttaaatatatacctaaag | Amplify tPGI1 with N-terminal ePTS1 tag |
| LN564 | (pTDH3)_EcBBE_F | caagaacttagtttcgaATAAACACACATAAACAAACAAAATGGAAAATAAGACACCGATTTTC | Amplify BBE with homology to pTDH3/tPGI1 |
| LN565 | (tPGI1)_EcBBE_R | ctttaatgttctttaggtatatatttaagagcgatttgttTTATATAACGACTTCTCCCCCG |  |
| PP118 | (LV3)_pPDC1_F | AGGAATACTCTGAATAAAACAACTTATATAATAAAAATGCacatgcgactgggt | Amplify genes from pGC997 with homology to integration sites |
| PP124 | (LV5)_tADH1_R | TGTTATAATTATTTTCTTATTTTGATGTAATATAAAGAGGgcatgccggtagag | Amplify MSH, P6H from pGC997 with homology to integration sites |
| PP127 | (LV5)_tPGI1_R | TGTTATAATTATTTTCTTATTTTGATGTAATATAAAGAGGggtatactggaggcttcat | Amplify MSH, P6H, and TNMT from pGC997 with homology to integration sites |
| LN1124 | SPS-FP | gcatcgtctcatcggtctcatatggaagaatctttctggattgtctcc | Amplify stylopine synthase for Golden Gate cloning |
| LN1125 | SPS-RP | atgccgtctcaggtctcaggatctaataaccgttaatagatggagaacct |  |
| LN1126 | CFS-FP | gcatcgtctcatcggtctcatatggaagaatctttctggttggttacc | Amplify cheilanthifoline synthase for Golden Gate cloning |
| LN1127 | CFS-RP | atgccgtctcaggtctcaggatctaatgagttcttggggtgatcttag |  |
| LN1128 | TNMT-FP | gcatcgtctcatcggtctcatatgggttcaatagatgaggtcaagaag | Amplify TNMT for Golden Gate cloning |
| LN1129 | TNMT-RP | atgccgtctcaggtctcaggatctacttcttcttgaaaagcagctgc |  |

Table S4. Synthetic gene list

*HsMAO-A* atggaaaatcaagaaaaggcatctattgcaggtcacatgtttgatgttgttgttataggtggaggaatttcaggtttgtcagctgctaaattgttaactgagtacggtgtttctgttttagtattagaggctagagacagagttggtggtagaacctatacaatcagaaacgaacatgtagactacgttgacgttggtggagcttatgttggtcctactcagaataggattttaaggttgtctaaggaattaggtattgaaacttataaagttaatgtttcagaaagattagtacagtatgttaaaggtaagacatatccatttagaggtgcctttccaccagtttggaatcctatcgcttacttggattacaataatttgtggagaaccattgataatatgggtaaggagatcccaactgacgcaccttgggaggcacagcatgcagataagtgggacaagatgaccatgaaggaattaatcgataaaatatgttggactaaaaccgcaaggagattcgcctatttatttgtaaatattaacgtaacttctgaacctcatgaagtttcagcattgtggttcttatggtatgtcaagcagtgtggtggtacaacaagaatattttcagttacaaacggtggtcaagaaagaaaattcgtcggtggttctggacaggtatctgaaagaattatggatttattaggtgatcaagtcaaattgaatcatccagttacccatgtagaccaatcttctgataacatcatcatagaaactttaaatcatgaacattatgagtgtaaatatgtcataaacgcaattcctccaaccttgactgccaaaatccattttagacctgaattacctgctgaaagaaaccaattaatccaaagattgccaatgggtgctgtaatcaaatgtatgatgtattacaaagaagctttctggaagaagaaagattactgtggttgcatgattatagaggatgaagatgctccaatttctatcaccttagacgatactaaacctgatggatctttacctgccataatgggattcatattggccagaaaggccgatagattggcaaagttacataaagaaattagaaagaagaaaatctgtgagttatatgctaaagtcttaggttctcaagaagccttgcatcctgtccattatgaagaaaagaactggtgtgaggaacaatattctggaggatgttacactgcttatttcccacctggaattatgacacaatacggaagagttatcaggcagccagtcggtagaattttctttgcaggaactgaaaccgctacaaaatggtctggttatatggaaggtgcagtagaggctggtgaaagggctgctagagaagttttaaatggattaggaaaagtcactgaaaaggatatttgggtacaggagcctgagtctaaagatgttcctgctgttgaaatcactcataccttttgggaaagaaacttgccatcagtctcaggtttgttgaaaatcattggtttctctacctcagtcaccgccttgggattcgttttgtataagtacaaattgttgccaagatca

*EcBBE (*from *Eschscholzia californica*)

atggaaaataagacaccgattttcttttcattatcaatcttcctatcattgctaaattgcgcgcttggcgggaatgatctattgtcatgcctaacgttcaatggtgtacgtaatcatactgtgtttagcgccgacagtgacagtgacttcaataggtttcttcaccttagtatacaaaaccctttgttccaaaactcactaatatccaagcctagtgctattatccttccggggagcaaagaggagttatccaacactataagatgtataagaaaaggttcctggaccataaggctgaggtcaggaggtcattcctacgaaggtctgtcatacaccagtgacacacccttcattcttatagacttaatgaatctaaatagggtaagtattgatttagaatcagaaacagcgtgggttgagagcggatcaactctaggagaattatattacgcgataacggaaagttcttcaaagcttggatttaccgcagggtggtgccctaccgtgggtacggggggtcacatctccggggggggctttggcatgatgagtagaaagtatgggctagctgctgacaatgtagtagatgcgatattaatcgacgcgaatggggctatcttagacaggcaagctatgggagaggacgtgttttgggccattagaggcggtggcggtggggtatggggagccatatacgcctggaaaataaagcttctacctgtccccgaaaaagtgaccgtattccgtgtaactaaaaacgtggcgatagacgaggcgacatcactgctacacaagtggcagttcgtcgctgaagaattggaagaggactttactctgtccgttctgggcggggcagatgagaagcaggtatggttgacgatgttaggctttcactttggactgaaaacagtagcaaagtcaacctttgatttacttttccctgaactaggacttgtagaggaggattaccttgaaatgtcctggggggaatccttcgcgtatctagctggattggaaaccgtgagccaacttaataatagattcctgaagtttgatgaaagggcattcaaaaccaaagttgatttaacgaaggagccgctgcctagcaaagcgttttatggtcttctggagagattatctaaagagccaaatggtttcatagctttgaacgggttcgggggacaaatgtcaaagatctcctcagacttcaccccgttccctcatcgttccggtacgaggttgatggtagaatacatcgtggcatggaatcaatcagaacaaaagaagaaaaccgagtttcttgactggttggaaaaagtttacgaatttatgaaaccgtttgtatctaagaatccccgtttgggatacgtcaaccatatagatttagatcttgggggaatagactgggggaataagacagtagttaacaatgcgatagagatcagtcgttcatggggtgagtcatattttttgtccaattacgagagattaatccgtgctaaaaccctaatagatcccaataatgtgtttaatcacccgcagtccatccccccgatggccaattttgattaccttgagaagacacttgggtccgacgggggagaagtcgttatataa
